# Genetic impacts on DNA methylation help elucidate regulatory genomic processes

**DOI:** 10.1101/2023.03.31.535045

**Authors:** Sergio Villicaña, Juan Castillo-Fernandez, Eilis Hannon, Colette Christiansen, Pei-Chien Tsai, Jane Maddock, Diana Kuh, Matthew Suderman, Christine Power, Caroline Relton, George Ploubidis, Andrew Wong, Rebecca Hardy, Alissa Goodman, Ken K. Ong, Jordana T. Bell

**Affiliations:** Department of Twin Research and Genetic Epidemiology, King’s College London, UK; University of Exeter Medical School, UK; MRC Unit for Lifelong Health and Ageing, Institute of Cardiovascular Science, University College London, UK; MRC Integrative Epidemiology Unit, University of Bristol, Bristol, UK; Population, Policy and Practice, UCL Great Ormond Street Institute of Child Health, University College London, UK; Centre for Longitudinal Studies, Institute of Education, University College London, UK; School of Sport, Exercise & Health Sciences, Loughborough University & UCL Social Research Institute, University College London, UK; MRC Epidemiology Unit and Department of Paediatrics, Wellcome Trust-MRC Institute of Metabolic Science, University of Cambridge School of Clinical Medicine, UK

**Keywords:** DNA methylation, heritability, GWAS, methylation quantitative trait loci, meQTL

## Abstract

Pinpointing genetic impacts on DNA methylation can improve our understanding of pathways that underlie gene regulation and disease risk. We report heritability and methylation quantitative trait locus (meQTL) analysis at 724,499 CpGs profiled with the Illumina Infinium MethylationEPIC array in 2,358 blood samples from three UK cohorts, with replication. Methylation levels at 34.2% of CpGs were affected by SNPs, and 98% of effects were *cis*-acting or within 1 Mbp of the tested CpG. Our results are consistent with meQTL analyses based on the former Illumina Infinium HumanMethylation450 array. Both meQTL SNPs and CpGs with meQTLs were overrepresented in enhancers, which have improved coverage on this platform compared to previous approaches. Co-localisation analyses across genetic effects on DNA methylation and 56 human traits identified 1,520 co-localisations across 1,325 unique CpGs and 34 phenotypes, including in disease-relevant genes, such *ICOSLG* (inflammatory bowel disease), and *USP1* and *DOCK7* (total cholesterol levels). Enrichment analysis of meQTLs and integration with expression QTLs gave insights into mechanisms underlying *cis*-meQTLs, for example through disruption of transcription factor binding sites for CTCF and SMC3, and *trans*-meQTLs, for example through regulating the expression of *ACD* and *SENP7* which can modulate DNA methylation at distal sites. Our findings improve the characterisation of the mechanisms underlying DNA methylation variability and are informative for prioritisation of GWAS variants for functional follow-ups. A results database and viewer are available online.

## 1 Introduction

DNA methylation is a major regulator of gene function, with important roles in development and over the life course [1–3]. In humans, DNA methylation and de-methylation occur predominantly at cytosine-guanine dinucleotides (CpG-sites) through the action of DNA methyltransferases and TET enzymes, respectively [4, 5]. The human methylome consists of a mosaic of regions exhibiting variable stability over time, including both longitudinally stable regions, as well as dynamic regions where changes can relate to ageing or reflect environmental exposures, such as smoking [6, 7].

Multiple studies have shown that genetic effects have considerable impacts on DNA methylation levels at specific CpGs. Family and twin-based estimates of narrow-sense heritability in DNA methylation levels in blood [8, 9] report a wide range from 0 to 1 heritability at individual CpGs profiled by the Illumina Infinium HumanMethylation450 array (450K). Most studies typically associate genetic variation at single nucleotide polymorphisms (SNPs), or methylation quantitative trait loci (meQTLs), to DNA methylation levels at a specific CpG. A large proportion of reported meQTLs are in close proximity to the tested CpG (usually within 1 Mbp, in *cis*), while long-range and inter-chromosomal associations (*trans*) only represent a small fraction of meQTL associations. A recent large-scale study in 27,750 European samples estimated that DNA methylation levels at up to 45% of CpGs in the blood 450K methylome are associated with meQTL SNPs [10], which are in turn more likely to be GWAS signals than expected by chance. Another recent analysis of 3,799 European and 3,195 South Asian samples further explored trans-ancestry effects, and confirmed multiple links between meQTLs and phenotype variation [11]. In addition to analyses based on blood, a variety of studies have also identified meQTL SNPs in different tissues, for instance in brain [12, 13], adipose [14] and buccal tissue samples [15].

The most extensively used profiling technology for human methylome analyses to date has been the Illumina 450K array, comprising approximately 480,000 probes [16], and the vast majority of meQTL reports are based on this platform. However, the 450K array has limited coverage outside of CpG islands (CGIs) and genic regions. The most recent Illumina methylation array, the Infinium MethylationEPIC BeadChip (EPIC), improves genomic coverage of enhancers which are key regulatory regions. The EPIC array assays 853,307 sites, adding 333,265 novel CpGs in enhancers to the near entire set of 450K CpGs [17]. Accordingly, there is a need for follow-up analyses to identify new genetic influences on DNA methylation levels profiled by the newer EPIC array. To date, only two studies have explored meQTLs on the EPIC array in blood, but both included relatively modest sample sizes (*n* = 156 *−* 1, 111) [18, 19] and did not consider both genome-wide *cis* and *trans*-meQTL effects.

Here, we report novel genome-wide meQTL analyses of DNA methylation profiles on the Illumina EPIC array, applying a meta-analysis across 2,358 samples from three UK population cohorts: TwinsUK [20], the MRC National Survey of Health and Development or 1946 British Birth Cohort (1946BC) [21], and the National Child Development Study or 1958 British Birth Cohort (1958BC) [22]. We characterised novel meQTLs specific to the Illumina EPIC array, and carried out genome annotation enrichments and meQTL integration with summary statistics from 54 GWASs and previously reported eQTLs. A database and viewer of results is available online.

## 2 Results

We explored genetic impacts on all CpGs profiled by the Illumina EPIC array by initially estimating twin-based narrow-sense heritability, and subsequently identifying common genetic variants associated with DNA methylation levels in *cis* and *trans* genome-wide. We independently analysed samples from each of the three UK cohorts (TwinsUK, 1946BC and 1958BC cohort) separately, and meta-analysed the results. Results are presented at a permutation-based false discovery rate (FDR). We validated a subset of our findings in an external meQTL catalogue from the GoDMC study [10], and replicated selected meQTLs in target regions using methylated DNA immunoprecipitation sequencing (MeDIP-seq) in an independent sample. Follow-up analyses included enrichment analyses within genomic annotations and ontologies, and co-localisation integrating meQTLs with previously reported eQTLs and summary statistics from 56 GWASs, as well as clumping SNPs based on linkage-disequilibrium (LD) (**Fig. 1**).

**Figure 1.**
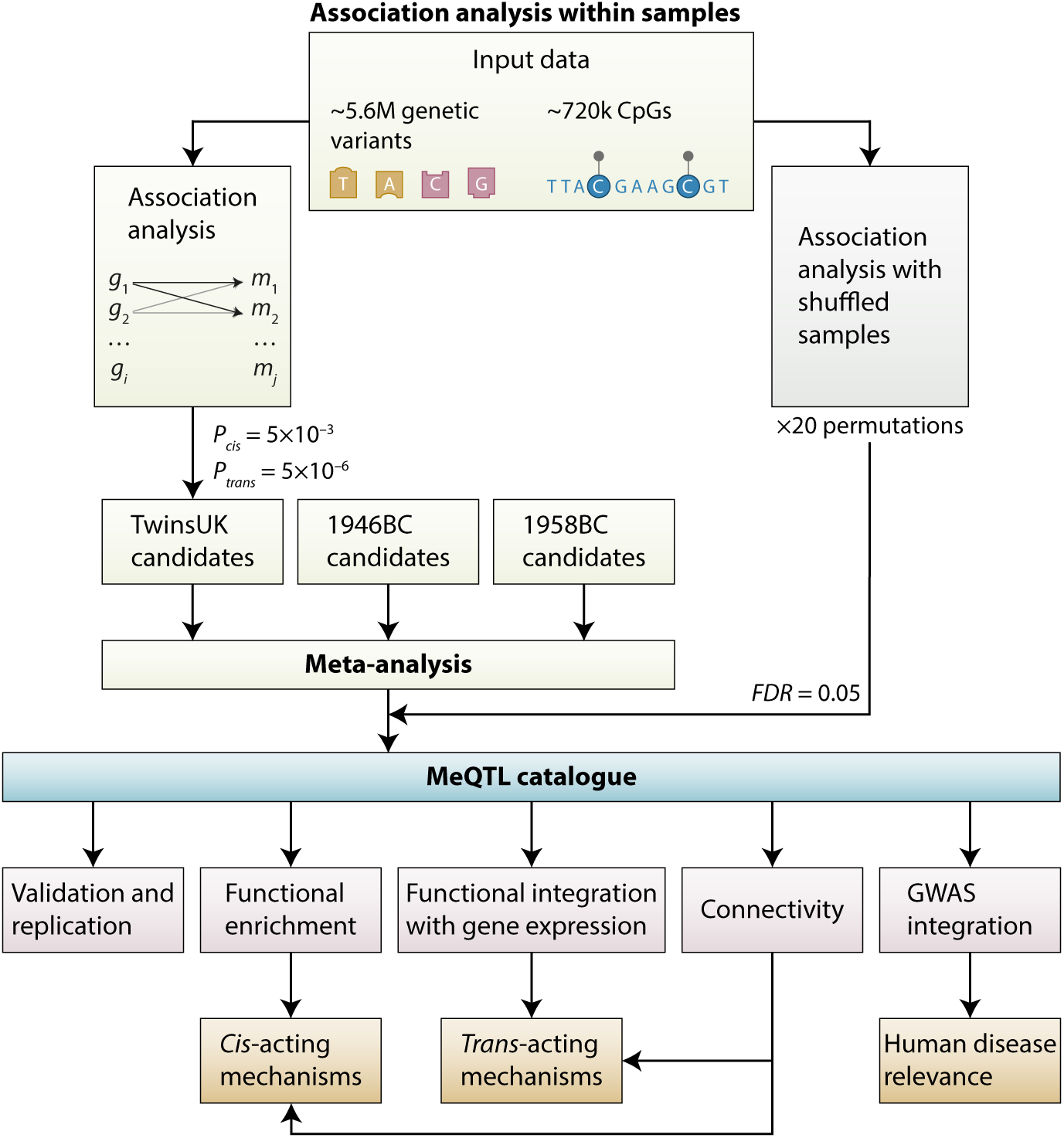
Study design. Genome-wide association analyses compared genotypes and DNA methylation levels profiled by the EPIC array. Each cohort sample was independently tested, and results were meta-analysed. Results are presented at a permutation-based false discovery rate (FDR). Follow-up analyses aimed to find evidence of underlying mechanisms and their relevance to human disease.

### 2.1 Heritability of the Illumina EPIC DNA methylome

We initially applied a classical twin study of 88 monozygotic (MZ) and 70 dizygotic (DZ) twin pairs from the TwinsUK cohort, to decompose the DNA methylation variance at each of 723,814 CpGs into additive genetic effects (*A*), common environmental effects (*C*) and nonshared environmental effects (*E*). The heritability distribution was zero-inflated (45.5% of sites have *A <* 0.01), and the maximum individual CpG heritability was 0.998 (cg21906335 in the promoter region of *ZNF155*; **Supplementary data 1**). Across all tested CpGs, the mean genome-wide narrow-sense heritability was *A* = 0.138 (*sd* = 0.198; median *A* = 0.037, IQR = 0.220) (**Fig. 2a**). When stratifying by genomic annotations, CpGs in enhancers tend to have overall greater heritability estimates (mean *A* = 0.179, *sd* = 0.217, 95% CI [0.178, 0.181]), for example, compared to promoters, which have one of the lowest heritability estimates (mean *A* = 0.106, *sd* = 0.179, 95% CI [0.105, 0.106]) (**Fig. 2b**). The improved representation of enhancer regions on the EPIC array may be reflected in a modestly greater mean heritability across novel EPIC-only sites (*A* = 0.142, *sd* = 0.198, *n* = 348, 091) than that across 450K legacy probes (*A* = 0.135, *sd* = 0.198, *n* = 375, 336; one-tailed *t*-test, *t*_(723,425)_ = 15.2, *P <* 2.2 *×* 10*^−^*^16^) (**Supplementary fig. 1a**). Overall, the heritability patterns of the genomic annotations are consistent between EPIC-only probes and 450K legacy probes (**Supplementary fig. 1b-c**), and are broadly in line with previously reported 450K heritability estimates across genomic annotations [9]. We also found that variable CpG sites (with methylation *β*-values *sd >* 0.025, see **Supplementary note**) tend to be the most heritable. For example, the average heritability of the most variable sites (*A* = 0.278) was double that estimated genome-wide (*A* = 0.138) and the zero-inflation rate was substantially lower (**Supplementary note**).

**Figure 2.**
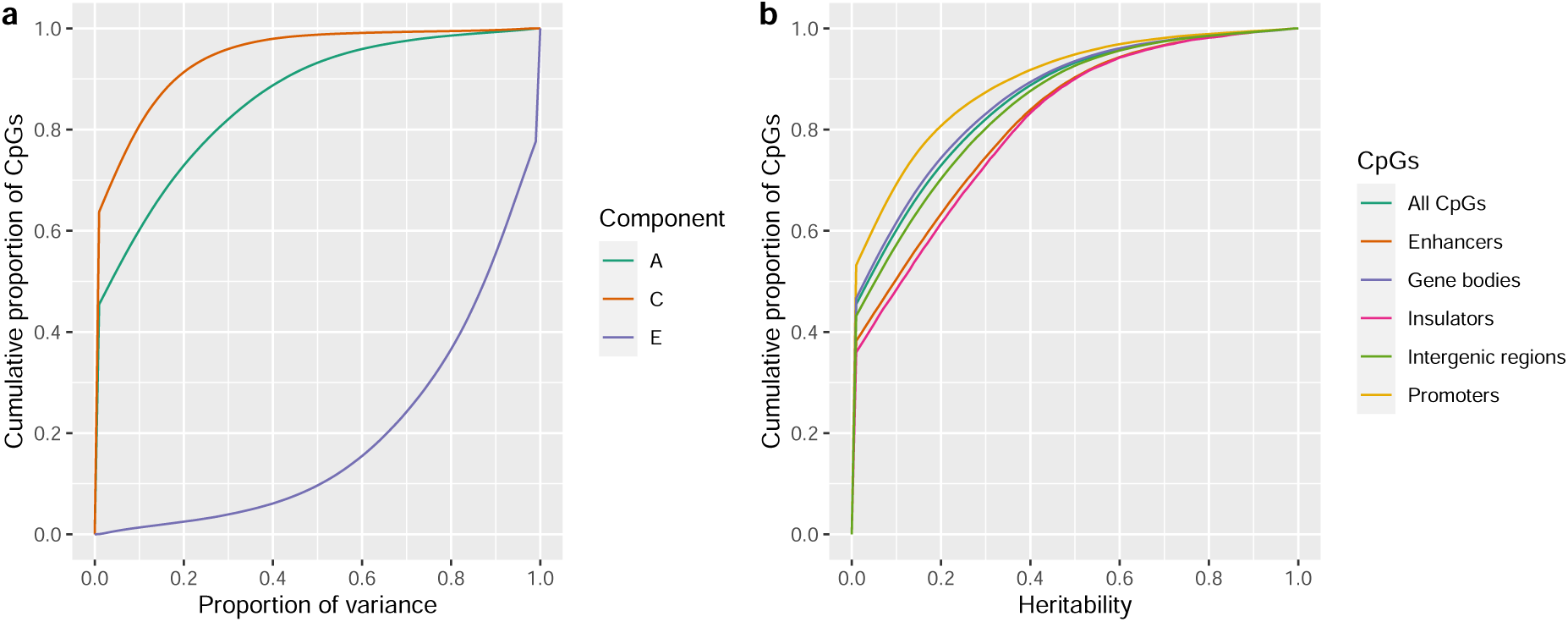
Proportion of variance of genome-wide DNA methylation levels attributed to genetic variation. Estimates for the 723,814 CpGs sites covered by the EPIC array after a classical twin study of 88 MZ and 70 DZ twin pairs from the TwinsUK cohort. (**a**) Cumulative proportion of variance components of the *ACE* model: variance explained by additive genetic effects, or heritability (*A*), common environmental effects (*C*) and nonshared environmental effects (*E*) (**b**) Cumulative proportion of heritability estimates by genomic annotations.

### 2.2 Common genetic variation has major impacts on the blood methylome

To identify specific genetic variants that impact the methylome, a meQTL analysis was carried out with a total of 2,358 whole blood samples across five datasets from three non-overlapping human cohort studies: TwinsUK, 1946BC and 1958BC (**Table 1**). Initially all SNP-CpG pairs were tested for association within each dataset. CpG and SNP associations within 1 Mbp (upstream and downstream) were considered to be in *cis*, and all others were considered to be in *trans* (**Supplementary table 1**). SNP-CpG associations that surpassed relaxed significance thresholds within each dataset were retained for meta-analysis, with a total of 189,202,234 unique candidate *cis* meQTL-CpG pairs (*P ≤* 5 *×* 10*^−^*^3^) and 100,814,822 *trans* pairs (*P ≤* 5 *×* 10*^−^*^6^). After meta-analysis we retained meQTL-CpG pairs where the strength of association surpassed FDR 5% (*P_cis_ ≤* 2.21 *×* 10*^−^*^4^, *P_trans_ ≤* 3.35 *×* 10*^−^*^9^), and where pairs were identified as candidates in more than one dataset with a consistent direction of effect.

**Table 1.** Summary characteristics for the five UK cohort sample sets.

| Cohort<sup>a</sup> | Sample size | Percentage females | Percentage smokers | Median age [Range] |
| --- | --- | --- | --- | --- |
| TwinsUK | 394 | 100 | 5.6 | 64.5 [42.4, 86.6] |
| 1946BC-99 | 1,348 | 52.3 | 24.2 | 53.4 [53, 54] |
| 1946BC-09 | 197 | 59.4 | 11.2 | 63.2 [60.3, 64.6] |
| 1958BC-1 | 183 | 50.8 | 21.3 | 45.1 [44.5, 45.8] |
| 1958BC-2 | 236 | 54.2 | 42.4 | 45.1 [44.3, 46.0] |
| **Total** | **2,358** | **60.9** | **21.5** | **53.5 [42.4, 86.6]** |

<sup>a</sup> 1946BC-99 and 1946BC-09 refers to independent samples from the 1946 birth cohort collected at two different time points and stratified to facilitate data handling. 1958BC-1 and 1958BC-2 refers to samples from the 1958 birth cohort processed in two different batches (see **Methods** for further details).

We identified 244,491 CpGs (33.7% of tested probes) to be under the influence of *cis*-meQTL SNPs, and 5,219 CpGs (0.7% of tested probes) to be influenced by *trans*-meQTL SNPs. Of these, 2,281 CpGs (0.9% of CpGs with *cis*-meQTL; 43.7% of CpGs with *trans*-meQTL) were influenced by at least one *cis* and one *trans* meQTL SNP simultaneously. There were 4,609,875 unique genetic variants identified as *cis*-meQTLs, and 240,866 identified as *trans*-meQTLs. Of these, 229,908 meQTLs were both *cis* and *trans*-meQTLs for CpGs at different sites. The meQTL SNPs and CpGs under genetic control altogether formed 39,110,128 *cis* and 805,319 *trans* SNP-CpGs pairs (**Fig. 3a**). We carried out sensitivity analysis by splitting the CpGs into two sets, 450K legacy probes and EPIC-specific probes, and repeating the meQTL discovery process, and found that the resulting proportions of meQTLs reported remained very similar (**Supplementary note**). The strength of the associations was stronger for *trans* SNP-CpG pairs than for *cis* SNP-CpG pairs, although this is an expected result due to the difference in *P*-value thresholds for *cis* and *trans* associations (**Supplementary note**). On the other hand, *trans* effects were more heterogeneous across samples compared to *cis* effects, and therefore the reported *trans* effects should be interpreted with caution. We estimated that, on average, a *cis*-meQTL explains 7.6% of the methylation variance of its associated CpG, while a *trans*-meQTL explains 11.5%, which is a significant difference (**Supplementary note**). CpGs with both *cis* and *trans* associations, and SNPs that act as both *cis* and *trans*-meQTLs, have associations with higher *R*^2^ estimates (**Supplementary note**). CpG-sites with meQTLs were evenly distributed across chromosomes according to number of genes per chromosome. However, this pattern was not observed for meQTL SNPs (**Supplementary note**).

**Figure 3.**
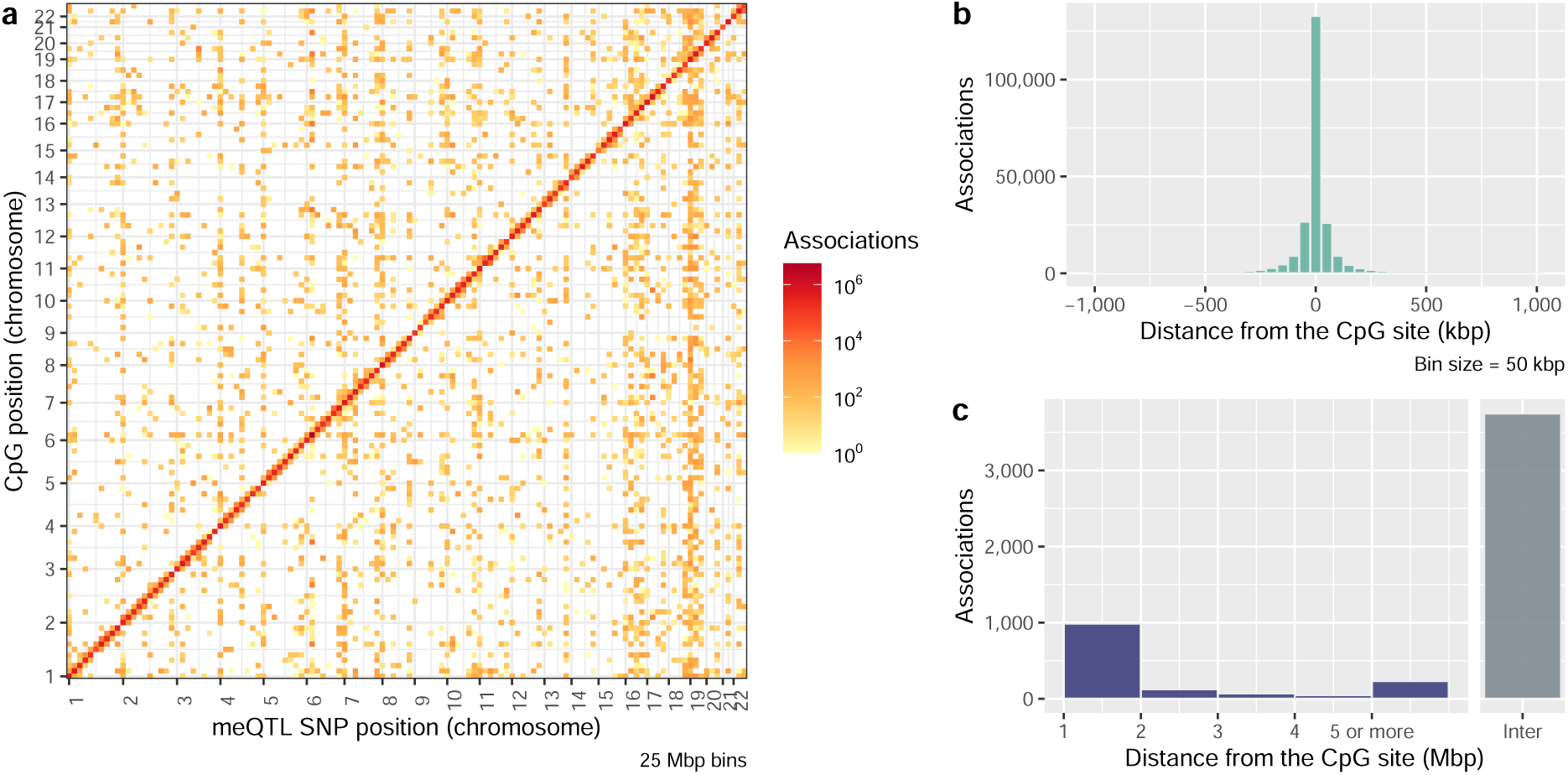
DNA methylation quantitative trait loci (meQTLs) for CpG sites genome-wide. Association analysis carried out between 724,499 CpGs vs. 6,361,063 SNPs. (**a**) Genomic distribution of meQTL associations at a significance level of FDR *<* 0.05. The *x*-axis corresponds to the position of the SNPs within in the 22 chromosomes and the *y*-axis to the position of the CpGs, with each pixel binning a range of 25 Mbps. The colour scale indicates the number of associations between those specific CpGs/SNPs locations, on logarithmic scale. (**b**) Histogram of distances between the CpGs and their most significant *cis*-meQTL SNPs. (**c**) Bar plot of absolute distances between the CpGs and their most significant *trans*-meQTL SNPs. Intra-chromosomal associations are shown in purple, and inter-chromosomal in grey.

*Cis*-meQTLs exhibited relatively short-range effects as expected [10, 18]. The median distance between each SNP *cis*-meQTLs and its target CpG was 20.5 kbp (interquartile range (IQR) = 65.5 kbp) if considering the most significant association (**Fig. 3b**), and 75.5 kbp (IQR = 165.5 kbp) if considering all significant associations (**Supplementary fig. 6a**). CpGs with *trans* associations have almost exclusively intra-chromosomal or inter-chromosomal meQTLs, and cases in which both types occur are rare (only 25 CpGs). For *trans* associations, 71.8% of the most significant associated SNPs per CpG are inter-chromosomal (**Fig. 3c**). When considering all *trans* associations the number of inter-chromosomal SNP-CpG pairs decreases to 45.4% (**Supplementary fig. 6b**).

We also explored evidence for cell type-specific meQTL effects. These analyses considered only *cis*-meQTLs effects specific to CD4^+^ T cells (mean ratio = 0.199, *sd* = 0.073 across samples) and monocytes (mean ratio = 0.05, *sd* = 0.026 across samples). We observed that 8.9% of all CpGs had *cis*-meQTL effects specific for either CD4^+^ T cells or monocytes (*P ≤* 2.21 *×* 10*^−^*^4^; see **Supplementary note** and **Supplementary table 2**). Our cell-specific results replicate a proportion of previously reported cell-specific meQTLs in CD4^+^ T cells (17.7%) and in monocytes (17.3%) [23]. Of the CpGs that had *cis*-meQTL SNPs in whole blood, 1.1% also showed evidence for cell-specific meQTL effects (*P ≤* 2.21 *×* 10*^−^*^4^), suggesting that the majority of genetic effects that we detect on CpGs in whole blood are stable across different blood cell types (**Supplementary note**).

Overall, meQTLs explain 14.2% of the variance in the DNA methylation heritability (*F*_(2,723,424)_ = 5.99 *×* 10^4^, *P <* 2 *×* 10*^−^*^16^) (**Supplementary fig. 8** and **Supplementary note**). CpGs without detected meQTLs have lower mean heritabilities (*A* = 0.085, *sd* = 0.146, *n* = 476, 192) compared to CpGs that have meQTLs. CpGs that have meQTLs can be split into three groups showing increasing mean heritabilities, from CpGs with only *cis*-meQTLs (*A* = 0.238, *sd* = 0.239, *n* = 242, 021), to CpGs with only *trans*-meQTLs (*A* = 0.295, *sd* = 0.251, *n* = 2, 933), and to CpGs with both *cis* and *tran*-meQTLs simultaneously (*A* = 0.435, *sd* = 0.259, *n* = 2, 281). Overall, CpGs with both *cis* and *trans*-meQTLs have the largest evidence for DNA methylation heritability.

### 2.3 Replication of novel EPIC-specific and 450K legacy CpGs with meQTLs

We pursued replication of meQTL effects at selected CpGs using previously published MeDIP-seq data in an independent sample of 2,319 individuals from the TwinsUK cohort. CpGs selected for replication included a subset of ten CpGs, which had the largest effect sizes (cg07143125, cg13904258, cg00918944, cg05808124), or with the largest number of meQTL SNP associations (cg25014118, cg00128506, cg18111489, cg16423305), or with meQTL SNPs that co-localised with GWAS signals (cg11024963, cg06162668). We replicated *cis*-meQTLs for 80% of the selected CpGs, and *trans*-meQTLs for 30% (**Supplementary table 3** and **Supplementary data 2**), after multiple testing correction and with a consistent direction of effect.

The EPIC array doubles the coverage of the 450K array. We observed that 51.2% of CpGs with *cis*-meQTLs (125,251 CpGs) and 37% of CpGs with *trans*-meQTLs (1,933 CpGs) are specific to the EPIC array. For the remaining 450K legacy CpGs with meQTLs, we validated our results by comparing them to the GoDMC database based on 32,851 blood samples [10]. Altogether, 97.0% of our 450K specific CpGs with meQTLs (in *cis* or *trans*) were also under genetic influence in the GoDMC dataset.

### 2.4 Genomic annotations of local and distal genetic effects show consistent enrichment in enhancers

We found overall contrasting patterns of genomic annotations for CpGs with *cis* and *trans*-meQTLs (**Fig. 4a** and **Supplementary table 6**). CpGs located in CpG islands (CGIs), promoters and transcription factor binding sites (TFBSs) are less likely to harbour *cis*-meQTLs (odds ratio (OR) *<* 1, FDR *≤* 0.05, two-tailed Fisher’s exact test), but are more likely to have *trans*-meQTLs (OR *>* 1, FDR *≤* 0.05). CpGs located in intergenic regions, enhancers and insulators are more likely to have both *cis* and *trans*-meQTLs. We also explored the overrepresentation of CpGs with meQTLs near to genes, and with respect to gene ontology (GO) terms related to molecular functions and biological processes (**Supplementary table 22**).

**Figure 4.**
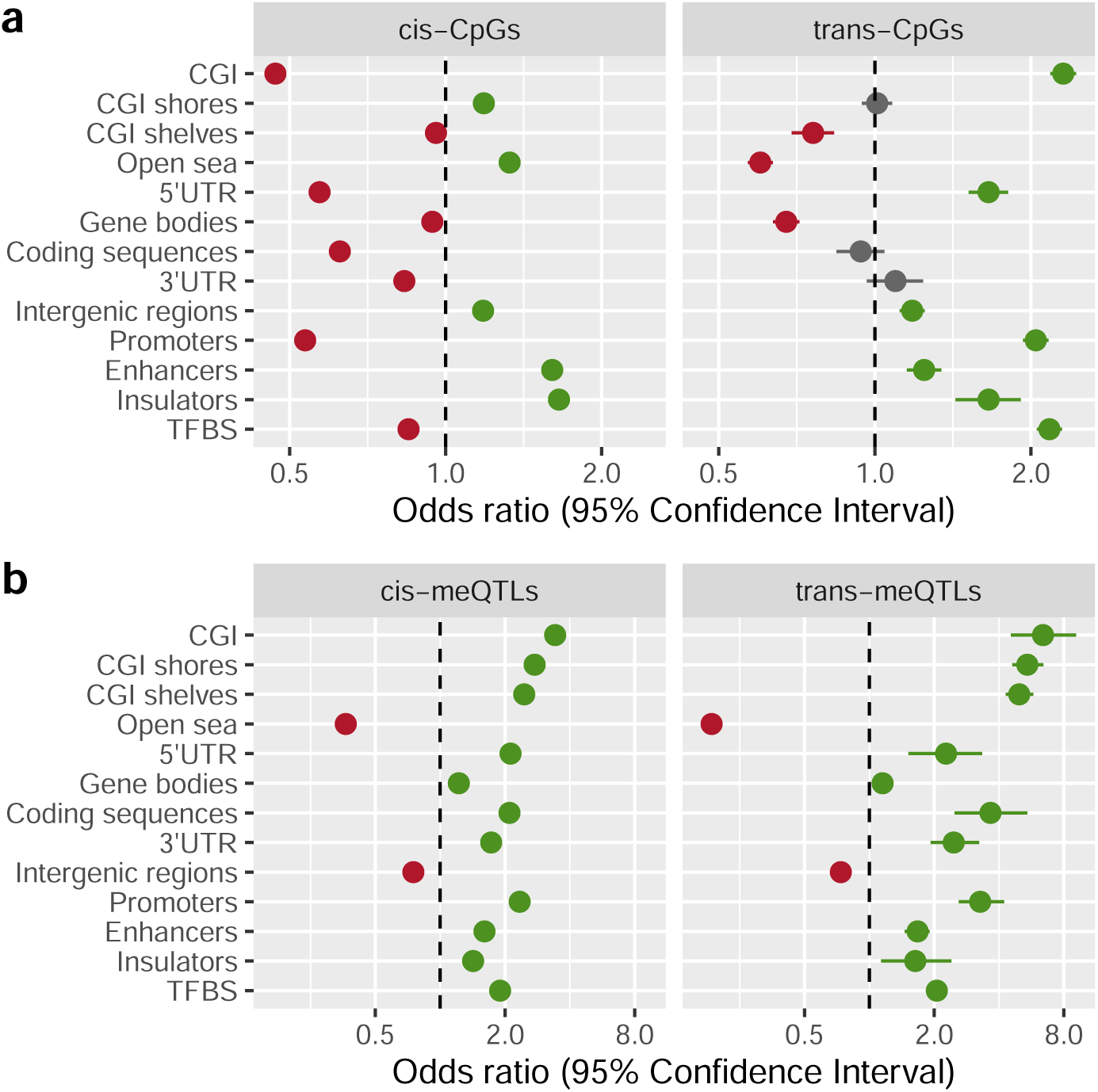
Enrichment in genomic annotations of meQTL SNPs and their CpGs. The *x*-axis indicates the odds ratio and its 95% confidence interval (in logarithmic scale) for (**a**) CpGs with meQTLs, or (**b**) meQTL SNPs, located within a specific genomic annotation. Significant enrichment is marked in green, depletion in red, and non-significant genomic annotations in grey.

We next explored enrichment or depletion of meQTL SNPs in different genomic annotations. To this end, we compared the proportions of the most significantly associated meQTL SNPs per CpG-site to the full panel of tested genetic variants in different annotations. Contrary to results observed for CpGs, we found a consistent pattern in the distribution of *cis* and the *trans*-meQTLs according to genomic category (**Supplementary fig. 9** and **Supplementary table 7**). Overall, coding regions, promoters, enhancers, insulators and TFBSs are over-represented for genetic variants that are meQTLs (OR *>* 1, FDR *≤* 0.05), either in *cis* or in *trans*. On the other hand, intergenic regions are under-represented for genetic variants that are meQTLs (OR *<* 1, FDR *≤* 0.05). The results remain consistent in sensitivity analyses taking as background reference subsamples of SNPs with a similar distribution of minor allele frequencies (MAF) and distance to target CpGs (**Fig. 4b** and **Supplementary table 8**). Therefore, unlike the genomic patterns observed for CpGs under genetic control, the genetic variants driving meQTL effects show similar genomic distributions for local and distal genetic effects.

The location of meQTL SNPs and CpGs helps to elucidate genetic mechanisms of methylome regulation. As previously proposed, TFBSs may play a critical role in *cis* associations, as a genetic variant could prevent protein binding and alter methylation of the surrounding loci [24–26] (**Fig. 5a**). The observed over-representation of meQTL SNPs in TFBSs, both for *cis* and *trans* results, supports this hypothesis (OR*_cis_* = 2.47, 95% CI*_cis_* [2.44, 2.50], OR*_trans_* = 1.92, 95% CI*_trans_* [1.74, 2.10]). We further explored this observation through enrichment analyses considering TFBS for sixteen specific transcription factors (TFs) of interest, previously identified as relevant for chromatin interactions and in the modulation of DNA methylation. These TFs included CTCF [27], ZNF143 [28] and EBF1 [29] (**Supplementary fig. 10**). The direction of the effect in all cases was consistent with the results from the overall TFBS enrichment analysis.

**Figure 5.**
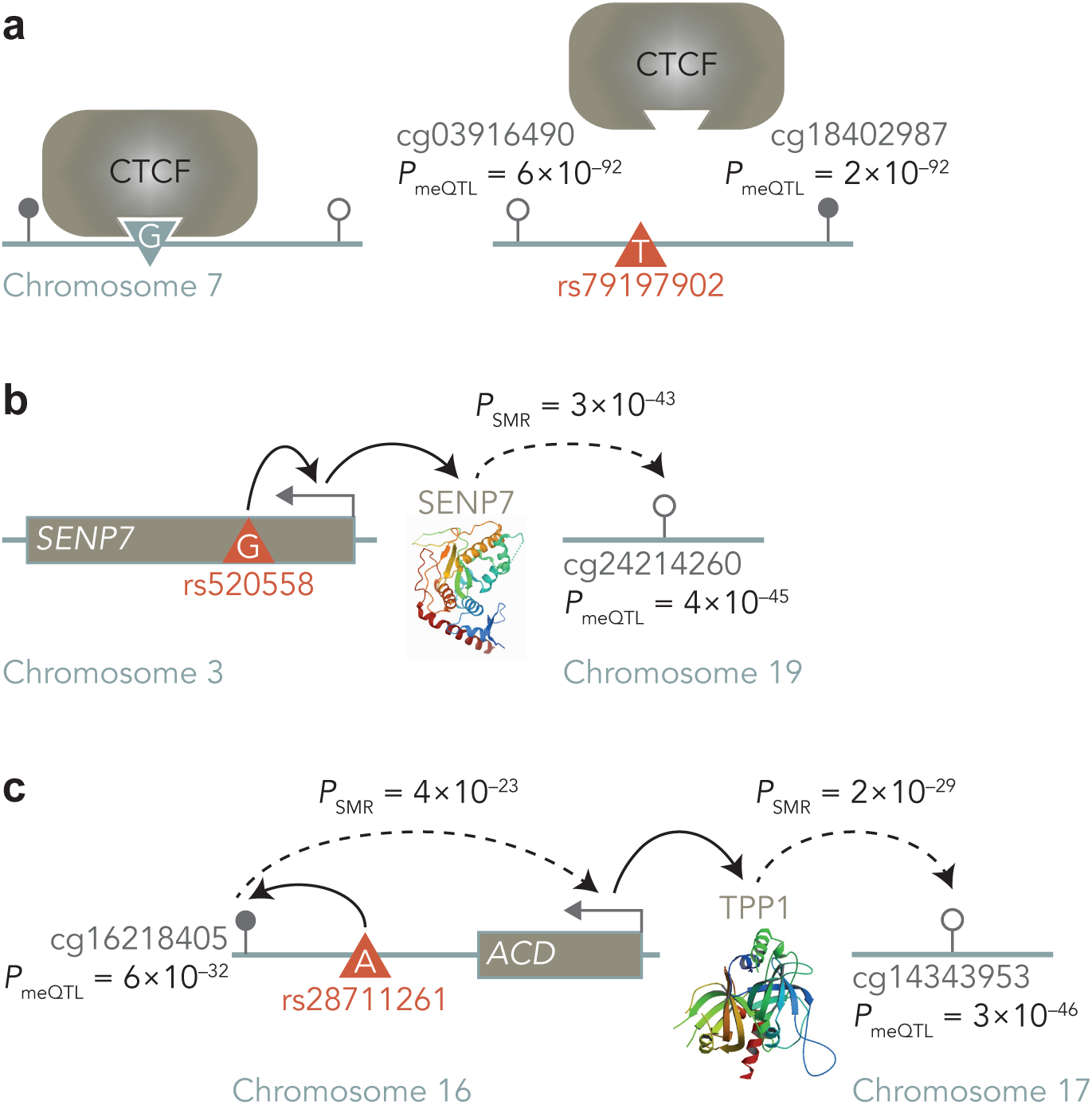
Underlying mechanisms of meQTL SNPs. (**a**) Example of a *cis*-meQTL mechanism. The disruption of a TFBS (e.g. CTCF binding site) by a genetic variant (rs79197902), leads to reduced affinity of the protein by its site, which changes local methylation (cg03916490, cg18402987). (**b**) Example of an ‘eQTL-mediation mechanism’ for *trans*-meQTLs. SNP rs520558 that is an eQTL for a gene involved in DNA methylation regulation (*SENP7*) indirectly affects distal CpG sites (cg24214260). Dashed lines represent associations for which there is suggestive, but not conclusive, evidence of directionality. (**c**) Example of a ‘*cis*-meQTL-mediation mechanism’ for *trans*-meQTLs. SNP rs28711261 is associated with a nearby CpG (cg16218405), which in turn is associated with a gene involved in DNA methylation regulation (*ACD* gene of TPP1), and indirectly affects distal CpG sites (cg14343953).

We inspected CpGs in enhancers in more detail, motivated by their targeted coverage on the EPIC array. A set of 39,450 CpGs with *cis*-meQTLs and 789 CpGs with *tran*-meQTLs were annotated to enhancers (strong and weak/poised enhancers, see **Methods**), based on ChromHMM annotations [30]. We find that the corresponding *cis*-meQTL genetic variants of CpGs in enhancers also tend to be in enhancer regions and TFBSs, when compared to the total set of meQTL SNPs (**Supplementary fig. 11** and **Supplementary tables 18–19**). This observation is not simply attributable to the genomic location of the associated CpGs in enhancers (**Supplementary note**). In short, we observe a clear enrichment of CpGs with meQTLs, and of meQTL SNPs, in enhancers.

### 2.5 Functional integration gives insights into long range genetic impacts on the methylome

To explore potential mechanisms underlying meQTL associations, we carried out several functional integration analyses.

First, we combined our meQTL findings with data from the eQTLGen Consortium [31], the most extensive eQTL resource to date, conducted on 31,684 blood samples from individuals from 37 cohorts of predominantly European ancestry. We used *cis*-eQTLs results for 19,250 genes, and applied Summary-based Mendelian Randomization (SMR) [32] to co-localise signals and infer putative pleiotropic or causal effects on DNA methylation and gene expression.

Overall, we observe robust evidence for co-localisation between *cis*-meQTLs and eQTLs, which is in line with previous findings [18, 33]. Analysis of *cis*-meQTLs identified 19,267 unique SNPs that co-localise with *cis*-eQTLs of 8,511 genes and with *cis*-meQTLs of 21,663 CpGs, resulting in 31,395 unique gene-CpG associations (*P*_SMR_ *≤* 9.82 *×* 10*^−^*^9^, *P*_HEIDI_ *>* 0.05) (**Supplementary data 3**). Altogether, 44.2% of expressed genes shared a genetic basis with DNA methylation, which is greater than previously reported [18, 34]. CpGs typically have shared genetic effects with a single gene (median = 1, IQR = 1). Site cg11024963 had the highest number of colocalization events (13 genes, including *DUS2*, *ZDHHC1*, *TPPP3* and *ECD4*) through the *cis*-meQTL rs8054034, successfully replicated in the MeDIP-seq dataset. Correspondingly, genes have shared genetic effects with a median of two CpGs (IQR = 2), and at most 88 CpGs, in the case of the *MSRA* gene. We observed an enrichment of CpGs with shared meQTL/eQTLs in genic and regulatory regions (**Supplementary fig. 12** and **Supplementary table 11**). The resulting SMR genes were related to immunological processes in GO analyses (**Supplementary table 23**). If we consider CpG-gene pairs with co-localised QTLs, we observe that methylation levels at the CpGs tend to be negatively correlated with the corresponding gene expression levels, regardless of the location of the CpG within the gene (**Supplementary fig. 13** and **Supplementary note**). In summary, we observe the largest to date shared genetic basis between local genetic impacts on DNA methylation and gene expression, suggesting presence of joint regulatory mechanisms.

SMR analysis with *trans*-meQTLs also identified a number of meQTLs and eQTLs co-localisation events. Altogether, 642 unique *trans*-meQTL SNPs co-localised with *cis*-eQTLs (1,520 co-localisation events), simultaneously affecting 709 CpGs and 782 genes (*P*_SMR_ *≤* 3.71 *×* 10*^−^*^7^, *P*_HEIDI_ *>* 0.05) (**Supplementary data 3**). A median of one CpG is associated per gene (IQR = 1) and one gene per CpG (IQR = 2). The results could reflect a scenario where genetic variants that influence the expression of genes involved in direct or indirect global epigenetic regulation are also *trans*-meQTLs (i.e. ‘eQTL-mediation mechanism’ from Villicaña & Bell [26], also proposed by Huan et al. [35]). The gene with the most associations to CpGs through co-localised QTLs was *SENP7* (19 CpGs in chromosome 19, one on chromosome 5 and one on chromosome 10; **Fig. 5b**). Our findings are in line with recent studies indicating that *SENP7* interacts with epigenetic regulators in the context of DNA repair [11, 36]. Within these CpG-gene pairs with shared *trans*-meQTLs/*cis*-eQTLs, we identified an enrichment of CpGs in enhancers and TFBSs (**Supplementary fig. 12** and **Supplementary table 11**). Furthermore, the genes are annotated to GO terms related to DNA-binding transcription repressor activity including predominantly zinc finger proteins, which are known to act as epigenetic regulators in different contexts [37–39] (**Supplementary table 23**). In addition, the results of GO enrichments also replicate findings from previous studies [11, 35].

The *cis*-meQTL to *cis*-eQTL co-localisation results also allow us to make inferences into mechanisms of distal genetic impacts on DNA methylation levels. We observed a significant enrichment of *trans*-meQTLs in the co-localised *cis*-meQTL to *cis*-eQTL SNPs, compared to the non-co-localised *cis*-meQTLs (OR = 3.73, 95% CI [3.59, 3.88]). In light of this, we then used these *trans*-meQTL SNPs (that co-localised with *cis*-meQTL and *cis*-eQTL) as instrumental variables in SMR to test for associations between the corresponding eQTL gene expression levels and DNA methylation levels of the corresponding CpGs in *trans*. We identified a total of 511 *trans*-associations through 279 SNPs (hereafter ‘multi-QTLs’), between 323 CpGs and 292 genes (*P*_SMR_ *≤* 9.82 *×* 10*^−^*^9^, *P*_HEIDI_ *>* 0.05) (**Supplementary data 3**). These results could reflect a genetic mechanism of *trans*-meQTL effects, where a *cis*-meQTL impacts nearby CpG sites. These CpG sites in turn may affect the expression of genes involved in epigenetic regulatory processes, and whose products affect the methylation of multiple distal sites (i.e. ‘*cis*-meQTL-mediation mechanism’ from [26]). The adrenocortical dysplasia homolog (*ACD)* gene fits this scenario (**Fig. 5c**). *ACD* has four eQTLs (rs28711261, rs9936153, rs12935253 and rs2059850989) that co-localised with *cis*-meQTLs of six CpGs, and *trans*-meQTLs of 16 CpGs on different chromosomes. *ACD* produces the TPP1 protein, which is part of the shelterin complex that maintains telomere length [40]. A correlation between DNA methylation patterns and telomere length has been reported previously [41, 42], although multiple mechanisms likely underlie these links given that condensation of telomeric chromatin by the shelterin complex does not primarily occur through DNA methylation [43].

We carried out two additional functional exploration analyses of meQTLs. First, we searched for meQTL-CpG associations that overlapped three-dimensional (3D) conformations of the genome, such as topologically associated domains (TADs). The rationale behind this analysis was that some intra-chromosomal *trans*-meQTLs may act as ‘long-range’ *cis*-meQTLs [11, 35] that TADs bring into physical proximity [26, 44]. We integrated our meQTL results with TADs predicted from multiple-tissue Hi-C experiments [45–49]. We found that 36.5% of CpGs with intra-chromosomal *trans*-meQTLs share the same TAD with their most associated meQTL. In comparison, 17.1% of CpGs with *cis*-meQTLs share the same TAD with their most associated meQTL. Furthermore, TADs containing intra-chromosomal *trans*-meQTL associations are significantly larger than TADs with *cis*-meQTL associations (mean TAD size*_cis_* = 1.2 Mbp; mean TAD size*_trans_* = 3.4 Mbp; *P ≤* 2.54 *×* 10*^−^*^13^), which supports our hypothesis that TADs may bring *trans*-meQTLs into physical proximity with their target CpG (**Supplementary note**). In summary, our results are consistent with the hypothesis that some intra-chromosomal *trans*-meQTLs may act as ‘long-range’ *cis*-meQTLs within TADs.

Second, we focused on GO analysis of *trans*-meQTLs that lie within coding regions to test for evidence that *trans*-meQTLs may alter the function of proteins such as TFs. Our motivation was that such SNPs may impact the binding affinity of the TFs, and therefore change DNA methylation levels of distal unoccupied binding sites. A total of 79 *trans*-meQTLs (1.8% of the 4,398 top *trans*-meQTLs) were annotated in coding regions of 168 protein-coding genes. We found enrichment in 37 GO terms relative to molecular functions and 182 terms for biological processes (**Supplementary table 24**). Of these, 11 corresponded to categories related to protein binding and 56 to regulation of biological processes. Therefore, these results support the hypothesis that *trans*-meQTLs may alter the function of proteins such as TFs that then impact DNA methylation levels at multiple genomic regions.

### 2.6 Highly connected CpGs and meQTLs

We calculated the effective number of meQTL SNP associations per CpG, discarding redundant SNPs due to LD. To this end, we merged all *cis*-meQTL SNPs and following LD clumping generated ‘*cis*-meQTL regions’, and repeated the process for *trans*-meQTLs (see **Supplementary note**). Overall, CpGs under genetic control tend to have few associations after LD clumping, with a median of two meQTLs in *cis* (IQR = 3) and one meQTL in *trans* (IQR = 1) per CpG. However, a subset of CpGs have a high number of clumped meQTLs, or are ‘highly regulated’ or connected. Specifically, such highly regulated CpGs include 1.4% of CpGs with *cis*-meQTLs that have over 13 clumped meQTLs, and 2.9% with *trans*-meQTLs with over 5 meQTL associations (thresholds correspond to *Q*_3_ + 3IQR). From the CpGs with both *cis* and *trans*-meQTLs (2,281 sites), 627 CpGs are highly regulated by either *cis*-meQTLs, *trans*-meQTLs or both.

Highly regulated CpGs with *cis*-meQTLs are overrepresented in genic and regulatory regions, such as enhancers, compared to other CpGs with *cis*-meQTLs (**Supplementary fig. 14**, **Supplementary tables 12** and **20**). In the case of highly regulated CpGs with *trans-*meQTLs, coding sequences are enriched, while promoters, TFBSs and intergenic regions are depleted. Moreover, 32 immune-related GO annotations are enriched for highly regulated CpGs in *cis* but not in *trans* (**Supplementary table 25**). The CpGs that have the most associations overall both *cis* and *trans* are the novel EPIC probes cg16423305 (42 *cis* and 21 *trans*-meQTLs), cg00128506 (48 *cis* and 13 *trans*-meQTLs) and cg25014118 (50 *cis* and 6 *trans*-meQTLs). All these CpGs are located on chromosomal region 8p23.1 near to or in genes *PRAG1*, *MFHAS1* and *XKR6*. Additionally, CpG cg00128506 is in an enhancer region, and in the binding site of transcription factors ELF1, USF2, IKZF2 and RAD51. We replicated 81% of cg00128506 *cis* and *trans-*meQTL associations in the MeDIP-seq dataset (**Supplementary table 3**).

We next considered the connectivity of meQTL SNPs. We observed a median of five unique *cis*-CpGs (IQR = 10) associated with each region of clumped meQTLs, and a median of one *trans*-CpG (IQR = 1). Highly connected meQTL clumped regions, or ‘key regulatory regions’, were defined as 4.4% (*cis*) and 7.8% (*trans*) of genetic regions associated with more than 42 *cis*-CpGs and 5 *trans*-CpGs, respectively (thresholds correspond to *Q*_3_ + 3IQR). A relatively large proportion of 71.9% of the meQTLs that act simultaneously in *cis* and *trans* (165,290 SNPs) are located in key regulatory regions in either *cis*, *trans* or both. MeQTLs located within key regulatory regions are enriched/depleted in the same genomic regions as the top meQTLs previously described (**Supplementary fig. 15**, **Supplementary table 13** and **21**).

Particularly noticeable among highly connected CpGs and genetic regions is the major histocompatibility complex (MHC) region, which is overrepresented with both CpGs with genetic effects and SNPs that are meQTL. This locus contains multiple highly regulated CpGs and key regulatory meQTL regions for *cis* and especially *trans* associations (**Supplementary note**). However, the very high genetic diversity and complexity of this genomic region necessitates further follow-ups with higher resolution genetic and epigenetic sequence datasets. Apart from the MHC region, other genomic regions with high level of CpG connectivity include the above-mentioned region on chromosome 8p23.1 (6,200,001– 12,700,000 region spanning many genes). For meQTL SNP-level connectivity other genomic region hotspots included chromosomes 17q25.3 (in *B3GNTL1*) and 21q22.3 (in multiple genes) for *cis* associations, and 19p13.2 (*ZNF* gene family) and 7p22.3 (*MAD1L1*) for *trans* associations.

We also compared the number of clumped meQTLs per CpGs in enhancer regions. We detected a small but significant increase in the mean number of *cis*-meQTL associations for CpGs in enhancers (3.14 meQTLs per CpG outside of enhancers, compared to 3.58 in enhancers; two-tailed *t*-test, unequal variances, *t*_(52,211)_ = 23.7, *P <* 2.2 *×* 10*^−^*^16^), but no difference in the median number of associations (two *cis*-meQTLs for both categories). For *trans*-meQTLs, neither the mean (two-tailed *t*-test, unequal variance, *t*_(1,042.7)_ = 0.4, *P* = 0.69) or the median differed across these categories. Overall, the CpG sites with most genetic associations are found in enhancers, as confirmed by the enrichment observed of highly regulated CpGs.

### 2.7 The interplay between genetic variation, DNA methylation and complex traits

We used our meQTL findings to identify co-localisations between our *cis*-meQTLs SNPs and GWAS SNPs from 56 common human complex traits grouped in seven phenotypic categories (**Supplementary table 26**).

After the Bonferroni correction for 186,817 CpG-sites tested and the seven phenotypic classes, we identified 1,520 associations through co-localisation between 1,325 unique CpGs and 34 traits, involving 1,180 unique *cis*-meQTLs (*P*_SMR_ *≤* 3.82 *×* 10*^−^*^8^, *P*_HEIDI_ *>* 0.05) (**Supplementary tables 26–27** and **Supplementary data 4**). Height was the phenotype with the most GWAS signals co-localised with meQTLs (501 CpG-sites). ‘Growth and ageing’ (which includes height) was the phenotypic class with most co-localisations and 598 unique CpG-sites. The CpGs with most associations were cg06162668 and cg27288595, with six traits each. CpG cg06162668 is in an intergenic region in chromosome 2, and was associated with obesity and metabolic disease phenotypes through SNP rs7561317. The association between cg06162668 and rs7561317 was replicated in the MeDIP-seq dataset. Site cg27288595 was also associated with obesity and metabolic disease, along with growth and ageing, and is located in the *ZBTB38*. This gene encodes a zinc finger that binds to DNA methylation sites and acts a transcriptional repressor [37]. Overall, CpGs with GWAS co-localisations are enriched in CGIs, coding and regulatory regions, compared with other CpGs with meQTLs (**Supplementary fig. 18** and **Supplementary table 17**).

The strongest associations were between total cholesterol levels with cg17260184 and cg27123834, annotated upstream of the transcription starting site of *USP1* and *DOCK7*, respectively. *USP1* encodes a deubiquitinase which regulates the cellular response to DNA damage [50]. *DOCK7* —primarily involved in axon formation and neurogenesis—also overlaps the gene encoding angiopoietin-like protein (*ANGPTL3)* that regulates plasma lipid levels [51, 52]. The associated genetic variants rs2131925 and rs12136083, respectively, are in non-coding regions. To our knowledge, the function of these two variants has not been characterised.

Another example of note is the observed association between inflammatory bowel disease (IBD) and cg19297788 (*β*_SMR_ = *−*0.41, *P*_SMR_ = 1.44 *×* 10*^−^*^12^, *P*_HEIDI_ = 0.06), a CpG in a weak enhancer region of chromosome 21 (**Fig. 6**). The CpG also falls within three TFBSs for TCF12, EBF1 and RUNX3 and was not previously covered by the 450K array. We found evidence of association with both conditions comprised by IBD, ulcerative colitis (*β*_SMR_ = *−*0.39, *P*_SMR_ = 9.27 *×* 10*^−^*^9^, *P*_HEIDI_ = 0.11) and Crohn’s disease (*β*_SMR_ = *−*0.47, *P*_SMR_ = 3.11 *×* 10*^−^*^10^, *P*_HEIDI_ = 0.08). This locus is surrounded by five genes, including *ICOSLG*, a coding gene for a ligand of the T-cell surface receptor ICOS. This gene has been identified in previous studies as a risk locus for IBD [53–55], where the interaction between ICOS/ICOSLG in IBD and decreased expression of *ICOSL* can affect IBD risk [54]. However, it was unclear how genetic variants in the locus lead to the change in gene expression of *ICOSL*. According to our results, IBD and CpG-site cg19297788 share the common genetic variant rs2876932 (chr21:45,618,536).

**Figure 6.**
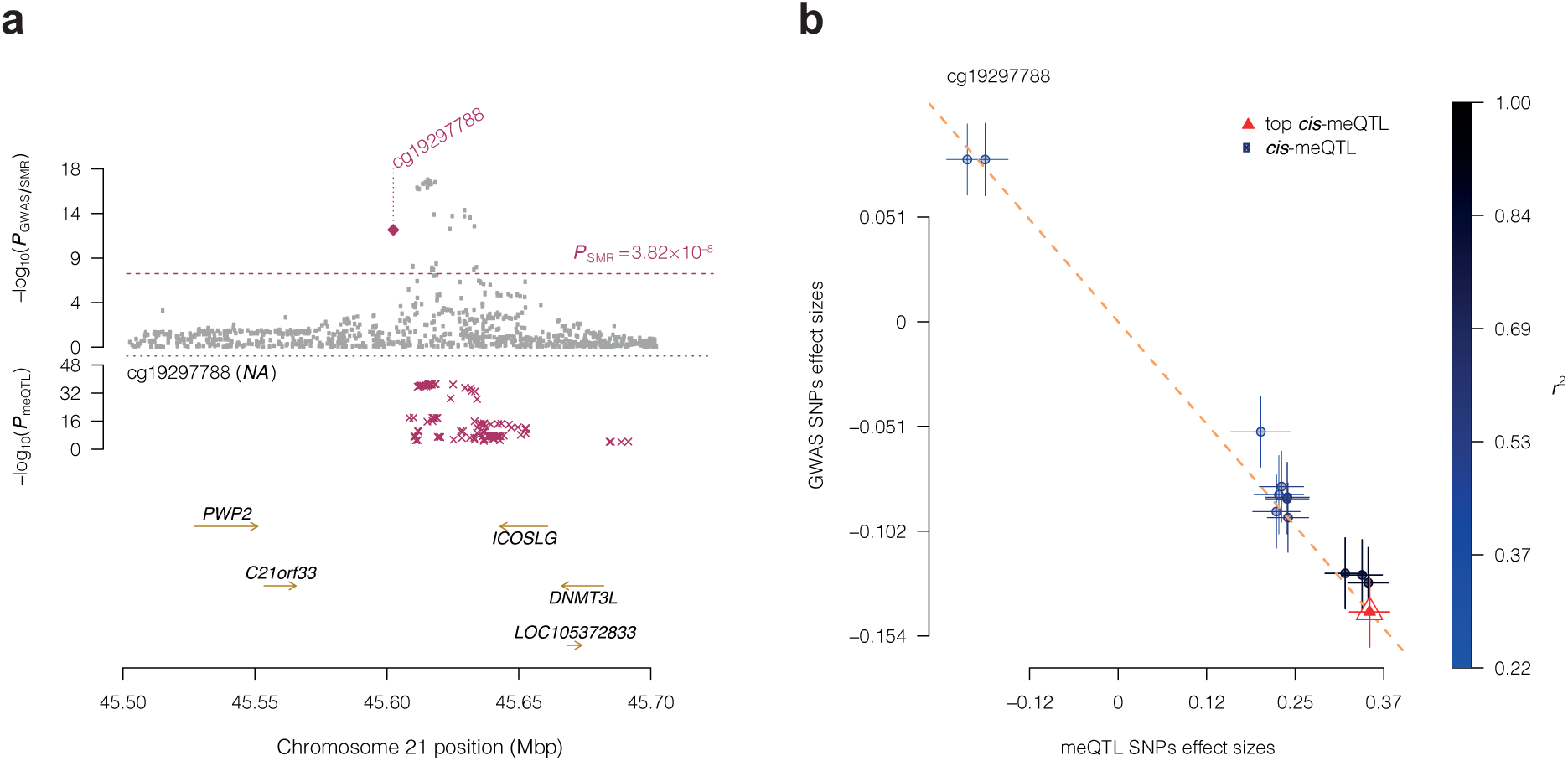
Association between IBD and DNA methylation at site cg19297788. (**a**) Locus association plot. The grey dots represent the *P*-values of the SNPs from the IBD GWAS^53^, the violet diamond the *P*-value of the SMR test, and the violet crosses the *P*-values of the meQTLs of cg19297788. (**b**) Effect sizes of IBD GWAS SNPs vs. effect sizes of meQTLs of cg19297788, for SNPs used in the HEIDI test. The slope of the dashed line represents the *β*_SMR_ estimate at the co-localised SNP. Error bars represent standard errors of estimated SNP effects. SNPs in LD with the top co-localised meQTL are expected to have a consistent effect under the causality/pleiotropy scenario.

Another example of note includes the associations observed for CpG cg17459721 and phenotypes for waist (*β*_SMR_ = *−*0.11, *P*_SMR_ = 1.73 *×* 10*^−^*^12^, *P*_HEIDI_ = 0.98) and hip circumference (*β*_SMR_ = *−*0.11, *P*_SMR_ = 3.59 *×* 10*^−^*^12^, *P*_HEIDI_ = 0.98), through rs7187776 on chromosome 16p11.2. Previous GWAS have described this region in the context of body mass index and body fat distribution [56, 57], but the mechanisms of action remain unclear. Here we identified from the SMR with gene expression that this meQTL SNP co-localises with eQTLs of the *TUFM* and *SPNS1* genes, and *trans*-meQTL for cg03969070 (chromosome 1). The latter CpG is in the promoter of *STK40*, involved in the glycogen metabolism (GO:0005977), among other biological processes. Therefore, we hypothesise that the action of rs7187776 is through a *cis*-meQTL-mediation mechanism.

Altogether these examples of integrative analyses highlight connections between target genetic variants and DNA methylation at multiple CpGs, gene expression at several genes, and a number of complex metabolic traits and diseases. These novel links provide functional insights into mechanisms of action for specific GWAS variants in selected human phenotypes.

## 3 Discussion

We investigated the impact of genetic variation on DNA methylation levels at genomic regions profiled by the Illumina Infinium MethylationEPIC BeadChip in three UK cohort populations. To our knowledge, previous meQTL studies have not yet explored both *cis* and *trans*-meQTLs across the genome on the new EPIC array in a large number of samples in blood. The increased coverage of the array, especially in intergenic regions such as enhancers, provides novel insights into the genetic regulation of DNA methylation, with downstream impacts into the regulation of gene expression and human complex traits.

We estimated that more than 33% of the EPIC methylome is under genetic control, the majority of which is in *cis*. Our *cis* results are in line with previous studies on the Illumina EPIC and 450K, in terms of proportion of sites, distance to target, allele frequency, and genomic annotations [10, 11, 19, 35]. The proportion of *trans* signals that we detected is somewhat lower than previous studies [10, 18, 35], although this likely in part reflects power as our two-stage meta-analysis approach may reduce power to detect *trans* associations. Specifically, before filtering associations in at least two cohorts, the detected *trans* associations were 10-fold greater compared to the final list (compared to *cis*, only 2-fold higher). This likely represents lower reproducibility of *trans* signals, which may be more likely subject to cohort specific differences, batch effects, or may potentially represent biological factors. Therefore, the reported *trans*-meQTL results should be interpreted with caution and validated in future studies. Furthermore and consistent with previous studies [11, 35], we also observe evidence that intra-chromosomal *trans*-meQTLs are likely to be ‘long-range’ *cis*-meQTLs, as the vast majority are located within 5 Mbp from the target CpG and a proportion fall within TADs. Lastly, in line with recent large-scale findings from the blood 450K meQTLs [10], our results confirm that SNPs and CpGs that exhibit both *cis* and *trans* associations, are highly reproducible, appear to have large effects, and exhibit high connectivity with other genetic variants and CpGs. Our results also highlight multiple highly connected genomic regions of interest, both putative key regulatory regions of SNPs, and regions containing highly-regulated CpGs. These connectivity results improve our understanding of specific mechanisms of genetic regulation on the epigenome, transcriptome, and human phenotypes.

We estimated the proportion of variance explained by meQTLs, both in relation to variability of DNA methylation at each CpG and to methylation heritability. Although the mean values genome-wide appear relatively low, there are cases of CpGs where genetic factors explain close to 100% of CpG DNA methylation variance. A similar trend is observed in terms of the number of meQTLs per CpG, where there are few CpGs with a large number of associations after LD clumping. These extreme cases, instead of being seen as exceptions, can be further explored in future to better understand the underlying mechanisms, evolutionary selection, and epistatic and environmental interactions of meQTL.

We integrated our meQTL results with large-scale blood eQTL results, as well as with GWAS findings from 56 human phenotypes. Altogether, these integrative analyses highlighted sets of shared genetic impacts that allow us to make two key inferences. The first one through eQTL integration gives insights into specific mechanisms of long-range genetic impacts on DNA methylation, highlighting multiple examples consistent with two hypothesized mechanisms. Second, through integration of GWAS findings with meQTLs, and in cases with eQTLs, we highlight multiple examples of specific putative mechanisms underlying GWAS genetic impacts on human phenotypes. Our work is consistent with and extends previous efforts, both disease-specific [58, 59] and multi-trait [18, 34], that integrate different molecular data at the genome-wide level to provide new insights into disease processes and biological pathways.

One of the main strengths of our study is that the sample used is representative and age-homogenous of a well-characterised nationwide population. Limitations include, first, analyses were restricted to whole blood samples. Although blood cell heterogeneity was taken into account, the estimated cell proportions are relatively low resolution. We undertook cell-specific analysis and observed that the majority of whole blood meQTLs do not show evidence for cell-specific effects. However, we did not comprehensively explore cell-population specific meQTL effects and restricted our analysis to two cell types with modest to moderate proportions in our whole blood data. Second, we did not include conditional analyses and therefore the number of independent meQTLs per CpGs remains unknown. Third, we carried out validation of all legacy 450K signals in the GoDMC dataset, and pursued replication at targeted novel EPIC-specific sites. The resolution of MeDIP-seq methylation data (500 bp) is lower compared to EPIC data and therefore presents a more qualitative replication approach. Fourth, limitations to eQTL and meQTL integration include the assumption of shared genetic impacts, although the effects may be coincidental. Fifth, several limitations of the SMR approach include no explicit test for causal impacts, a limited selection of 56 phenotypes considered, and differences in power across phenotypes because different GWASs have differing samples sizes and therefore power. Sixth, we cannot rule out that our sample has a selection bias, with an overrepresentation of healthy participants able to give blood samples and information on health. Finally, our findings relate to middle-aged and older adults. Although there is evidence to suggest that the meQTL effects are stable across the life course [60], further studies should confirm whether the associations described here are valid in other age groups. In this same line, we cannot extrapolate our observations to other ethnic groups.

In summary, we present a novel large-scale DNA methylation quantitative trait locus analysis in blood samples from three UK cohorts profiled on the Illumina EPIC array. The results identify novel genetic impacts on DNA methylation levels across the genome, and integrative analyses with gene expression and GWAS findings give insights into mechanisms underlying genetic regulation of human functional and phenotypic variability.

## 4 Methods

### 4.1 Study cohorts

We analysed data collected from 2,478 samples across three different UK population cohorts, of which 2,358 samples passed quality control assessment and are included in the analyses in this manuscript. TwinsUK [20] (post-QC *n* = 394, from 236 unique families) is the UK’s most comprehensive and detailed registry of adult monozygotic and dizygotic twins. The MRC National Survey of Health and Development (NSHD), or 1946 British birth cohort (1946BC) [21] (post-QC *n* = 1, 545), is the longest-running birth cohort in the UK, with data about individuals born during one week in March 1946. The National Child Development Study (NCDS), or 1958 British birth cohort (1958BC) [22] (post-QC *n* = 419), surveys individuals born during the same week in March 1958. The 1946BC data contained samples of individuals at either age *≈* 53 or *≈* 63, and therefore, we stratified the cohort in two age-based groups to facilitate data handling, referred to as 1946BC-99 (*n* = 1, 348) and 1946BC-09 (*n* = 197). The 1958BC samples were processed in two different batches and also stratified into 1958BC-1 (*n* = 183) and 1958BC-2 (*n* = 236). Local research ethics committees granted ethical approval of the study, and all participants provided written informed consent.

### 4.2 Genotyping and imputation

DNA was extracted from whole blood samples and genotyping was carried out with a combination of platforms across studies (**Supplementary note**). Quality control of raw genotype data from each of the five samples was carried out separately in PLINK [61], and steps included filtering out low-frequency and rare variants (minor allele frequency, MAF *<* 0.01), with a Hardy-Weinberg equilibrium *P <* 1 *×* 10*^−^*^6^ or missingness rate *>* 3%. We also removed samples with more than 5% of missing data. We imputed genotypes with the 1000 Genomes reference panel phase 3 version 5 [62] in the Michigan Imputation Server [63] and again filtered the resulting variants using a threshold for MAF *>* 0.05 and *r*^2^ *>* 0.8. For the present study we used the genome assembly GRCh37/hg19 [64] for reporting the genomic positions. The final set was of 6,361,063 unique genetic variants in at least one of the sample sets (**Supplementary table 1** with CpG-sites per cohort).

### 4.3 DNA methylation profiling and data processing

DNA was bisulfite-converted using the EZ DNA methylation kit (Zymo Research). DNA methylation levels were profiled with the Infinium MethylationEPIC BeadChip (EPIC) at a site-specific resolution, and raw intensities signals were obtained. Altogether five cohort samples were profiled, and detailed description of profiling and DNA methylation data initial processing is provided in **Supplementary note**.

Briefly, raw intensities signals were processed (separately for each sample set) with the ENmix package [65] in the R environment [66] and converted into Illumina *β*-values (ratio of methylation at each CpG-site) for downstream analysis. Background correction was performed using the Exponential-Normal mixture distribution (ENmix) method, dye-bias correction was performed using the Regression on Logarithm of Internal Control probes (RELIC) method [67], and probe design bias adjustment was performed implementing the Regression on Correlated Probes (RCP) method [68]. Filtering included exclusion of probes with missingness rates *>* 5% (detection *P >* 1 *×* 10*^−^*^6^) and exclusion of samples with missing methylation data at *>* 5% CpG (detection *P >* 1 *×* 10*^−^*^6^) and with no genotyping data. Additionally, we filtered out probes with a polymorphism with MAF *>* 0.05 in the interrogated CpG or the extension base (in case of type II probes), using the UK10K haplotype reference panel, plus the recommended list of masked probes published by Zhou et al. [69]. After data normalisation, we retained 724,499 unique CpGs in the autosomes across the sample sets (**Supplementary table 1** with CpG-sites per cohort). For the analysis, the number of samples with DNA methylation and genotyping data was 2,358 (see **Table 1** for the final sample size of each cohort).

### 4.4 DNA methylation data adjustment

DNA methylation profiles are cell type-dependent, and cell composition is a major confounding variable in methylation studies in tissues with cellular heterogeneity, such as whole blood [70]. We estimated the cell composition for monocytes, granulocytes, plasmablasts and immune cells (natural killer, näıve CD8^+^, CD4^+^, and joined CD8^+^/CD28^−^/CD45RA cells), using the regression calibration approach proposed by Houseman et al. [70] and implemented in the R package minfi [71].

To ensure normality and reduce the impact of confounders in the analyses, we applied a rank-based inverse normal transformation (INT) to the DNA methylation *β*-values and fitted a linear mixed-effects model (LMM) with covariates with the lme4 package [72]. We specified as fixed effects the variables sex (only for the birth cohorts), blood cell proportions, smoking and age (only for TwinsUK), and as random effects the technical covariates plate ID and position on the chip (as well as family ID and zygosity for TwinsUK). The residuals of this model were used for downstream analyses.

### 4.5 Heritability estimation

We used a classical twin design to estimate the narrow-sense heritability (*h*^2^) of DNA methylation at a CpG-level for TwinsUK data, with the OpenMx package [73] in R. After removing singletons, we kept 70 MZ twin pairs and 88 MZ twin pairs from the cohort. We used adjusted residuals of *β*-values (without the correction of family ID and zygosity) of the 723,814 CpG-sites available in the cohort. We applied structural equations and maximum likelihood estimation to decompose the variance proportion at each CpG-site in additive genetic (*A*), shared environment (*C*) and unique environment plus residual (*E*) components. The *h*^2^ corresponds to the proportion of phenotypic variance attributed to additive genetic effects (*A* component). We discarded CpGs where the model had critical optimization failures, keeping estimations for 723,427 CpGs. We compared the mean heritability between novel EPIC-only sites and the 450K legacy probes using a one-tailed *t*-test assuming equal variance.

### 4.6 Genome-wide association of DNA methylation

MeQTL analysis was performed in two stages. In the discovery phase, we identified candidate associations per sample. We fitted a linear regression between all possible pairs of SNPs and CpG-sites, with the genotype variant as the explanatory variable—coded as doses of the alternative allele (0, 1 or 2)—and adjusted *β*-values for the CpG as the response. In total, 3.4 billion of *cis* pairs and 4.7 trillion of *trans* pairs were tested across the five cohort samples. SNPs separated by no more than 1 Mbp from the tested CpG were considered *cis*, and the remaining *trans*. In the discovery step we applied a liberal *P*-value to keep the associations for further analysis, specifically, *P ≤* 5 *×* 10*^−^*^3^ for *cis* and *P ≤* 5 *×* 10*^−^*^6^ for *trans* associations. The discovery step was performed in Matrix eQTL [74] implemented in R.

The second stage was a meta-analysis with the summary statistics of the subset of candidate associations kept from the discovery phase. As some of the sample sizes of the cohorts are substantially different, which impacts the variance of the estimated coefficients, we accounted for this heterogeneity in a random-effects inverse-variance weighted meta-analysis, using the open-source software GWAMA [75].

To account for multiple testing, we estimated the false discovery rate (FDR) with a permutation approach. Briefly, for each of the cohorts, we shuffled the labels of the individual samples for the methylation profiles (maintaining the family structure in TwinsUK), and association tests on the permuted data were carried out as before in Matrix eQTL and meta-analysed in GWAMA. A total of twenty permutations were performed overall, and the resulted *P*-values formed our null distribution. Then, we calculated the FDR as described in Hastie et al. [76], with the proportion of associations in the null distribution over the associations in the observed real data. SNP-CpG pairs were reported as significant meQTLs if they had an FDR *≤* 0.05 (*P ≤* 2.21 *×* 10*^−^*^4^ for *cis*, *P ≤* 3.35 *×* 10*^−^*^9^ for *trans*). Lastly, we only report those associations that were observed in two or more of the five sample sets and with the same direction of effect. As a sensitivity analysis, we also estimated the threshold *P*-values by dividing the EPIC CpGs into two sets (legacy 450K probes only and the novel EPIC-only probes). All the details are available in the **Supplementary note**.

We replicated our results with the GoDMC meQTL catalogue [10]. We selected from our list of CpGs probes, those that were included in the 450K array and that were in the GoDMC study. We considered CpGs to replicate if they were also reported to be under genetic influence in the GoDMC study, with the same or with different SNP as that identified in our study.

For the integration of meQTL and heritability results, we fit a linear regression with the *A* estimate for each CpG as the dependent variable, and the categorical variables indicating the presence or absence of *cis*- or *trans*-meQTLs as independent.

### 4.7 Cell type-specific meQTLs

In addition to identifying whole blood meQTLs, we also explored evidence for blood cell-specific meQTL effects. To this end we considered DNA methylation-based estimates of blood cell proportion for each sample cohort (**Supplementary fig. 7a**), and focused on CD4^+^ T cells and monocytes. In cell-type specific analyses for CD4^+^ T cells, we first adjusted DNA methylation levels for covariates as described in the whole blood meQTL analysis, but did not include estimated proportion of CD4^+^ T cells as a covariate. We then fitted a linear model to estimate *cis*-meQTLs in Matrix eQTL. We considered the genetic variant, the proportion of CD4^+^ T cells, and the interaction term between these as predictors. For each cohort sample, we kept all associations where the interaction term surpassed *P ≤* 5 *×* 10*^−^*^3^. We then meta-analysed the summary statistics of the interaction terms in a random-effects model using GWAMA, and filtered associations observed in two or more sample sets with the same direction of effect. We used a similar process to estimate cell-specific meQTL effects for monocytes.

### 4.8 Linkage disequilibrium (LD)-based clumping of meQTLs

To account for LD structure among the identified meQTLs, we carried our LD clumping of the meQTL SNPs, performed separately for *cis* and *trans* meQTLs. Here, we kept the genetic variant with more associated CpGs as representative for each LD block—to ensure that all clumps were consistent across all CpGs. LD clumping was performed using PLINK with LD threshold of *r*^2^ *>* 0.1 (calculated using all the samples in this study) within a window of 2 Mbp. Finally, as the representative SNP of each clump may not be the one associated with a given CpG, we used the most significant meQTL per CpG and per clump.

### 4.9 MeDIP-seq data

For meQTL replication of novel EPIC probes we used previously published methylation data profiled with methylated DNA immunoprecipitation sequencing (MeDIP-seq) in TwinsUK blood samples [77, 78]. We excluded individuals from the current study, resulting in a final independent sample of 2,319 participants (from 1,632 unique families, 93.5% females, median age 55, age range 16–82) from the TwinsUK cohort, with whole blood methylomes profiled using MeDIP-seq.

MeDIP-seq of whole blood samples was performed as previously described [78]. Briefly, following DNA fragmentation through sonication, sequencing libraries were prepared using Illumina’s DNA Sample Prep kit for single-end sequencing. The anti-5mC antibody (Diagenode) was used for immunoprecipitation and MeDIP was validated by quantitative polymerase chain reaction. Captured DNA was purified and amplified with adaptor-mediated PCR, and fragments of size 200–500 bp were selected by gel excision and QC assessed by Agilent BioAnalyzer. Sequencing was carried out on the Illumina platform. Sequencing data were aligned using BWA [79] using build GRCh37/hg19 and a mapping quality score of Q10, and QC steps included FastQC, removal of duplicate reads, and SAMTools [80] QC. MeDIP-seq data quantification into methylation levels was carried out using MEDIPS v1.0 [81] reads per million (RPM), and further QC was carried out in R including batch effects inspection. The average high quality BWA aligned reads were *≈* 16.8 million per sample. Processed MeDIP-seq methylation data for analysis were quantified in genomic windows (bins) of 500 bp (250 bp overlap) with RPM scores.

We selected ten novel EPIC CpGs to replicate based on the number of associations, the strength of association, effect sizes, and the co-localisation of their meQTLs resulting after SMR. MeDIP-seq methylation levels in each bin were transformed with the rank-based INT and adjusted for covariates (sex, age, family and zygosity) in an LMM. We excluded bins with evidence of methylation association with smoking (*P <* 0.05). Finally, we excluded bins with methylation data in less than 1,160 samples. The final set of CpGs and the respective bins is listed in **Supplementary table 3**.

We performed the meQTL analysis in Matrix eQTL as described above. We considered associations to replicated if they exceeded a statistical threshold of *P <* 0.005 (Bonferroni correction for 10 CpGs at a significance level of 0.05) and with the same direction of effect as in the original EPIC meQTL analysis.

### 4.10 Functional annotations

We obtained different genomic annotations in BED file format through UCSC Table Browser as of August 31, 2020 [82]. We used annotations for CpG islands, RefSeq genes [83], chromatin state segmentation for the GM12878 cell line [30], and TFBSs for GM12878 cell line from ENCODE 3 [84]. Then we mapped the DNA methylation sites and genetic variants with the functional annotations.

Chromosome sizes and number of coding genes were retrieved from the reference genome GRCh37/hg19 and Ensembl 104 [85] databases. We considered the major histocompatibility complex (MHC) locus to span chromosome 6 base-pair positions 28,477,797 to 33,448,354 bp [64].

We incorporated 3D genomic annotations in the meQTL functional annotations by using previously published data from Hi-C experiments in lymphoblastoid cell line GM12878 [46, 47]. We also considered additional Hi-C data across multiple relevant cells and tissues, including in GM12878, in spleen [48] and thymus [49]. TADs from these datasets were the generated in and obtained from the 3D Genome Browser [45]. We estimated the percentage of CpGs where the target CpG and its most associated meQTL fell within the same TAD. This analysis was carried out for all CpGs with *cis*-meQTLs and for all CpGs with intra-chromosomal *trans*-meQTLs. The estimation was performed for the GM12878 Hi-C data alone, as well as for TADs estimated from multiple cells and tissues. Further details and results are provided in the **Supplementary note**.

### 4.11 Enrichment analyses

Fisher’s exact tests were performed to investigate enrichment or depletion of CpGs/SNPs across genomic regions and functional annotations. We used a modified version of the R package LOLA [86] extending the default one-tailed to a two-tailed test, and incorporating the estimation of confidence intervals. The results for each independent analysis were corrected for multiple testing with the Benjamini–Hochberg procedure [87] to control the FDR.

For all enrichment tests on SNPs, we used the most significant meQTL per CpG (referred to as top meQTL). For some of the analyses on SNPs, we first generated a background set from a resampling method in order to obtain a collection of SNPs with equivalent distributions of MAFs and distances to the CpGs. To do this, we categorised the available SNPs according to their distance from the EPIC CpGs and their MAF. We took a random sample of SNPs with the same size as that of the set of interest (sample without replacement within each category, and with replacement across all the categories). Then, we did the enrichment analysis as described before, using the random sample as the background set. We repeated the process one thousand times and saved for each iteration the OR estimates. Finally, we obtained the mean OR of the annotations for the point estimate, and for the confidence intervals at *α* = 0.05, we used the 2.5% and 97.5% percentiles of resampling distribution of OR. All the enrichment tests results are presented specifying the set of CpGs/SNPs of interest and the background set used (**Supplementary tables 5–25**).

We carried out gene ontology (GO) [88] enrichment analyses with the R implementation of clusterProfiler [89]. This package uses a one-tailed Fisher’s exact test to find the overrepresentation of genes in molecular functions, cellular components or biological processes. *P*-values were corrected using Benjamini–Hochberg procedure, and redundancy was reduced in the enriched GO terms with the Wang method [90] using a similarity cut-off value of 0.7. GO enrichment analysis at a CpG level was performed with the GOmeth method available in the missMethyl package, which corrects for probe number and multi-gene bias [91].

### 4.12 Summary-based Mendelian Randomisation with gene expression data

We applied Summary-based Mendelian Randomization (SMR) analysis [32] between DNA methylation and gene expression. The SMR approach consists of two stages. First, it employs the most significant genetic variant associated with a CpG-site as an instrumental variable (IV) to test the association between the CpG and a phenotype through the two-step least-squares (2SLS) estimation. This association can be due to a causal relationship, horizontal pleiotropy, or LD. Under a causal/pleiotropic scenario, the estimations of the effect sizes of the DNA methylation on the expression levels are expected to be homogeneous when calculated with other SNPs in LD with the single causal variant. Therefore, for excluding spurious associations derived from LD, the second step is the heterogeneity in dependent instruments (HEIDI) test with up to twenty alternative SNPs for each CpG. A significant *P*-value in the HEIDI test is evidence of heterogeneity across the effects of the SNPs and, therefore, indicates that the phenotype and the CpG are associated with different causal variants in LD.

We used summary statistics from the eQTLGen Consortium [31], carried out on 31,684 blood samples from individuals from 37 cohorts (mostly of European ancestry). We stuck to the *cis*-eQTL results—pairs of SNPs and genes no more than 1 Mbp apart, considering the centre of the gene—available at 19,250 genes. We used SMR v1.03 with the default settings, with the European subset of 1000 Genomes phase 3 version 5 as reference panel.

In the first analysis, we only used the *cis*-meQTLs to find associations between genes and nearby CpGs. We tested a total of 5,092,588 pairs of CpGs–genes, setting a statistical threshold of *P*_SMR_ *≤* 9.82 *×* 10*^−^*^9^ (Bonferroni correction for a significance level of 0.05) and *P*_HEIDI_ *>* 0.05 to filter out associations due to LD.

The second SMR analysis was with the *trans*-meQTLs as IVs to test long-range associations through co-localised QTLs between all the CpGs and genes. We compared 134,698 CpG–gene pairs and established a significant threshold of *P*_SMR_ *≤* 3.71 *×* 10*^−^*^7^ (Bonferroni correction for a significance level of 0.05) and *P*_HEIDI_ *>* 0.05.

The final SMR eQTL analysis consisted of identifying associations between CpGs and distant genes through genetic variants that were also significant in the *cis* SMR analysis. For this, we made a list of targets SNP-CpG pairs (separated by more than 5 Mbp to exclude cases of long-range LD) to test for each gene and use the option *--extract-target-snp-probe* in SMR v1.03. We considered the same significance criterion as in the *cis*-meQTL–*cis*-eQTL co-localisation (*P*_SMR_ *≤* 9.82 *×* 10*^−^*^9^, *P*_HEIDI_ *>* 0.05).

### 4.13 Summary-based Mendelian Randomisation with GWAS data

We repeated the SMR approach to test for co-localisation of our significant *cis*-meQTLs with GWAS signals from 56 phenotypic traits, using summary statistics from previously published studies (details of each study in **Supplementary table 26**). We downloaded and prepared data, adding the chromosomal position of the variants using dbSNP 141 as reference [92] (where not annotated), and harmonising the ID format with that of the meQTLs. We used SMR v1.03 with the default settings, with the European subset of 1000 Genomes phase 3 version 5 as reference panel. We categorised *post hoc* the phenotypes into seven classes. For filtering co-localisations with sufficient statistical evidence, we set a threshold of *P*_SMR_ *≤* 3.82 *×* 10*^−^*^8^ after the Bonferroni adjustment of the significance level of 0.05 for the number of independent tests (186, 817 *×* 7 accounting the CpGs tested and the phenotypic classes), and a *P*_HEIDI_ *>* 0.05.

## 5 Data availability

An interactive web application with our results is available online (built using the R Shiny framework [93]), with summary statistics of top meQTLs and associations after LD clumping. **Supplementary data 1–4** present all the results of the analyses described here, including full summary statistics of significant meQTL associations.

Cohort-specific data availability details can be found in the **Supplementary note**.

## Acknowledgements

The work was supported by the UK Economic and Social Research Council (ESRC ES/N000404/1 to JTB), JPI-HDHL DIMENSION project funded in the UK via the Biotechnology and Biological Sciences Research Council (BBSRC BB/S020845/1 and BB/T019980/1 to JTB), and Mexican National Council of Science and Technology (CONACYT) doctoral fellowship (2019-000021-01EXTF-00323 to SV). Cohort-specific funding acknowledgements are included in the **Supplementary note**.

