## Supplementary note for "Genetic impacts on DNA methylation help elucidate regulatory genomic processes"

**3 Supplementary tables** **13**

**4 Supplementary figures** **17**

### 1 Supplementary methods and results

#### 1.1 Study cohorts and participants

##### 1.1.1 TwinsUK

TwinsUK is the largest twin registry in the UK with over 14,000 same-sex twins, established in 1992. Biological samples and data, and extensive clinical and questionnaire data are available in a large subset of cohort participants. The cohort and data collected have been previously described[1].

Epigenetic profiles were generated for 394 female research participants from TwinsUK at mean age 64.5. Subjects were not selected for any particular outcomes or exposures, but were selected to minimise missing data. The TwinsUK methylation data are uploaded on the ReShare UK Data Service, under Data collection id 853,526. Access to further individual-level data can be applied for through the cohort data access committee, see `https://twinsuk.ac.uk/resources-for-researchers/access-our-data/`.

Genotyping was done with a combination of Illumina arrays (HumanHap300, HumanHap610Q, 1M-Duo, and 1.2MDuo 1M).

All research participants have signed informed consent prior to taking part in any research activities. Ethical approval was granted by the National Research Ethics Service London-Westminster, the St Thomas' Research Ethics Committee (REC reference numbers: EC04/015 and 07/H0802/84).

##### 1.1.2 The MRC National Survey of Health and Development or 1946 British birth cohort

The 1946BC is the oldest of the British birth cohort studies, with data on health and life circumstances collected at multiple follow-ups from birth ($> 20$) on 5,362 men and women born in England, Scotland and Wales in March 1946. Epigenetic profiles were obtained from blood DNA samples from 1,545 1946BC participants at follow-ups at ages 53 and 60-64 years when intensive phenotype data collections were carried out, adding to a range and depth of existing prospective data across life. The cohort and the data collected have previously been described[2,3].

Epigenetic profiles were generated for two 1946BC samples. 1946BC-99, consisting of 1,348 subjects at mean age 53.4, was selected at random. 1946BC-09, consisting of 197 individuals at mean age 63.2 was selected to minimise data missingness for a wide range of exposures of the life course and phenotypes related to healthy ageing. Subjects were not selected for particular exposures or outcomes, or for extremes of phenotype distribution. DNA methylation profiles from 1946BC-99 and 1946BC-09 were profiled separately, but profiles were processed using the same procedure as described in the main manuscript (see Methods). The 1946BC-09 methylation dataset is available in the public domain. The

1946BC-09 methylation dataset access is through https://doi.org/10.5522/NSHD/S202. Access to further individual-level data can be applied for through the cohort data access committee, see http://www.nshd.mrc.ac.uk/data.

Genotyping was done with the MetaboChip custom genotyping array.

Ethical approval was granted by the Central Manchester Research Ethics Committee (07/H1008/168 and 07/H1008/245) and the Scotland A Research Ethics Committee (08/MRE00/12).

##### 1.1.3 The National Child Development Study or 1958 British birth cohort

The 1958BC is the second oldest of the British birth cohort studies. The initial sample of 17,415 individuals (8,411 females), consisting of all babies born in Great Britain in a single week in 1958, have had multiple follow-ups providing high quality prospective data on social, biological, physical, and psychological phenotypes at every sweep. This cohort and the blood samples taken have been described previously[4,5].

Epigenetic profiles were generated for two 1958BC samples. 1958BC-1, consisting of 183 subjects, was selected to minimise data missingness for a wide range of exposures of the life course and phenotypes related to healthy ageing. 1958B-1 subjects were not selected for particular exposures or outcomes, or for extremes of phenotype distribution. On the other hand, 1958BC-2 consisted of 236 subjects selected for extremes of child and adulthood adversity exposures. Briefly, the sample was selected according to a combinatorial exposure design for exposure either to low/high socio-economic position, childhood abuse or not, prenatal smoking or not, and bullying or not[6,7]. DNA methylation profiles from 1958BC-1 and 1958BC-2 were profiled separately, but profiles were processed using the same procedure as described in the main manuscript (see Methods). The 1958BC-1 and 1958BC-2 methylation datasets are available in the public domain. The methylation data access is through https://doi.org/10.5255/UKDA-SN-5594-2. Access to further individual-level data can be applied for through the cohort data access committee, see https://beta.ukdataservice.ac.uk/datacatalogue/series/series?id=2000032.

Genotyping was done with the Illumina Infinium Immunochip genotyping array.

Ethical approval was granted by the London Central REC (14/LO/0097, 12/LO/2010 and 08/H0718/29) and by South East MREC (01/1/44). This covered consent for the collection of blood samples for health research. Biosamples for the sweep are held at University of Bristol and this has ethical approval as a tissue bank under application 09/H1010/12 from North-West Haydock NRES committee.

#### 1.2 Heritability of variable CpG sites

We evaluated if variability of DNA methylation $\beta$-values influences the heritability estimation. We carried out a Spearman's test between the heritabilities and *sd* of the

$\beta$-values at each CpG site, and computed $P$-values via asymptotic $t$ approximation in R. The correlation between the variables is $\rho = 0.51$ , which is significatively greater than 0 ($S = 3.08 \times 10^{16}$, $P < 2.2 \times 10^{-16}$).

We filtered out CpG sites with low variability in methylation $\beta$-values across samples, using as cut-off value $sd < 0.025$, and repeated the genome-wide heritability estimations. A total of 238,988 CpGs (33% out of total sites) were included after the filtering. The distribution of the heritabilities changed, increasing the estimates for the mean ($A = 0.278$, $sd = 0.236$) and median ($A = 0.251$, IQR = 0.356), and reducing the zero-inflation rate (19.3% of sites with $A < 0.01$) (Supplementary fig. 2).

#### 1.3 MeQTL effect sizes

We compared the effect sizes of the *cis* and *trans*-meQTLs. We obtained the absolute value of the point estimate of the association coefficients and applied the inverse-variance weighted average. We performed a two-tailed weighted $t$-test, assuming unequal variance between the two groups (*cis* and *trans*), with the package weights in R[8].

We observed a significative difference of 0.14 units ($t_{(831,027)} = -772.4$, $P < 5 \times 10^{-5}$) between the mean effect size of *cis* and *trans* meQTL associations ($|\beta_{cis}| = 0.33$, $|\beta_{trans}| = 0.47$) (Supplementary fig. 3).

#### 1.4 Proportion of variance explained by meQTL effects

For the proportion of CpG methylation variance explained by individual meQTLs, we used as a rough estimate the coefficient of determination $R^2$ calculated from the largest sample (i.e. 1946BC-99, 1,348 samples). We had an $R^2$ estimate for 96.5% of *cis* associations (37,722,855) and for 99.3% of *trans* (799,986). The mean $R^2$ of *cis*-meQTLs is 0.076 ($sd = 0.094$) and median of 0.042 (IQR = 0.062, range [0.006, 0.887]). *Trans*-meQTLs have a mean $R^2$ of 0.115 ($sd = 0.084$) and a median of 0.091 (IQR = 0.073, range [0.015–0.811]) (Supplementary fig. 4). The difference in mean of the variance explained by the *cis*-meQTLs vs. *trans*-meQTLs is significant (two-tailed $t$-test, unequal variance assumed, $t_{(843,133)} = -410.3$, $P < 2.2 \times 10^{-16}$). We confirmed this result with a Mann-Whitney test as a sensitivity analysis (see Sensitivity analyses and Supplementary table 4).

Additionally, we compared the $R^2$ estimates of CpGs that have simultaneous *cis* and *trans* effects with those CpGs that only have one type of effect. We took the largest $R^2$ value as representative for each CpG. We found that SNP effects explained a higher proportion of the methylation variance of those CpGs that have both *cis* and *trans* associations at the same time (mean $R^2_{cis+trans} = 0.238$; mean $R^2_{cis/trans \text{ only}} = 0.115$; two-tailed $t$-test, unequal variance assumed, $t_{(2,296.6)} = 31.4$, $P < 2.2 \times 10^{-16}$). We confirmed this result with a Mann-Whitney test as a sensitivity analysis (see Sensitivity analyses and Supplementary table 4).

We followed the same approach to compare SNPs that are simultaneously *cis* and *trans*-meQTLs against all other meQTL SNPs. The results show that, on average, the former account for a higher proportion of methylation variance (mean $R^2_{cis+trans} = 0.307$; mean $R^2_{cis/trans\text{ only}} = 0.159$; two-tailed $t$-test, unequal variance assumed, $t_{(246,688)} = 387.8$, $P < 2.2 \times 10^{-16}$).

The association between site cg11670269 (chr6:37,588,605) and the *cis*-meQTL rs2179782 (chr6:37,588,563) has the highest $R^2$, with the SNP accounting for 88.7% of the variance of the methylation. Both the SNP and the CpG are in an intergenic region, at a TFBS. The *trans*-meQTL rs4783662 (chr16:68,719,649) associated with cg04657470 (chr2:198,365,150) has an $R^2$ of 0.811, being the long-range SNP that explains the highest proportion of methylation variance of a CpG. The variant rs4783662 is within an intron of the cadherin gene *CDH3*, whereas cg04657470 is in a locus annotated in a TFBS for multiple proteins, an active promoter and an exon of genes *HSPE1* and *MOB4*.

#### 1.5 Chromosomal distribution of CpGs and meQTL SNPs

A linear regression was used to assess if the number of CpGs or meQTL regions (after LD-clumping) was proportional to the number of genes per chromosome. The number of coding genes (obtained from Ensembl version 104[9]) was the independent variable in the models, and the dependent variable the number of unique CpGs or clumped meQTL regions with associations. We fit the regression for *cis* and *trans* separately.

We observed that CpG-sites with meQTLs are spread across chromosomes according to the number of genes in each chromosome, showing a strong correlation for *cis* ($F_{(1,\ 20)} = 49.68$, $P = 7.79 \times 10^{-7}$, $R^2 = 0.713$) and weak for *trans* ($F_{(1,\ 20)} = 7.62$, $P = 0.012$, $R^2 = 0.276$) (Supplementary fig. 5a). An exception is chromosome 6, with a large amount of CpGs under *cis* and *trans* genetic regulation. Specifically, MHC locus contains 4,307 CpGs from the 17,328 CpGs with *cis*-meQTLs in chromosome 6 (24.9%), and 568 out of 886 with *trans* (64.1%). On the other hand, chromosome 19 has a depletion of *cis*-CpGs, and *trans*-CpGs to a lesser extent.

Then, we repeated the same analysis for the LD-clumped regions that have meQTLs (see LD clumping section below). The chromosomal distribution as a function of the number of genes is less clear (*cis*: $F_{(1,\ 20)} = 15.02$, $P = 9.42 \times 10^{-4}$, $R^2 = 0.429$; *trans*: $F_{(1,\ 20)} = 21.17$, $P = 1.73 \times 10^{-4}$, $R^2 = 0.514$) (Supplementary fig. 5b). However, the enrichment of meQTL regions in chromosome 6 and the depletion on chromosome 19 (in *cis*) is consistent. Again, the MHC region overlaps a large number of clumped regions—3.1% and 22.8% of the *cis* and *trans*-meQTL clumps, respectively, of the total regions in chromosome 6.

#### 1.6 Cell type-specific meQTLs

We explored meQTL effects that act in a cell-specific manner. Overall, 8.9% of CpGs had meQTL effects specific for either CD4$^+$ T cells or monocytes (at $P \leq 2.21 \times 10^{-4}$; Supplementary table 2). The most significant interaction for CD4$^+$ T cells was the rs35958405 insertion (chr1:31,684,451) acting as a meQTL for cg18621232 ($\beta = 3.77$, $P = 3.42 \times 10^{-71}$), located in an intronic region of *NKAIN1* gene that codes for a protein interacting with the Na/K-ATPase (Supplementary fig. 7b). In the case of monocytes, the largest effect was from rs8075532 (chr17:72,617,027) on the site cg09648424, located upstream of the immune receptor gene *CD300E* (Supplementary fig. 7c).

The number signals detected across the entire EPIC panel is consistent with previous work [10]. Specifically, 8.9% of CpGs had meQTL effects specific for either CD4$^+$ T cells or monocytes (at $P \leq 2.21 \times 10^{-4}$). We compared our results with BLUEPRINT cell type-specific meQTLs [10], and 17.7% of our total CpGs replicate their results for CD4$^+$ T cells effects ($P \leq 2.21 \times 10^{-4}$, 2,434 out of 13,768 CpGs considering only 450K probes). For monocytes, 17.3% of our total CpGs replicate BLUEPRINT results ($P \leq 2.21 \times 10^{-4}$, 3,497 out of 20,169 CpGs considering only 450K probes).

If we only consider CpGs that have *cis*-meQTLs in whole blood, then we identify 70,321 *cis*-meQTL-CpG associations ($P \leq 2.21 \times 10^{-4}$) with cell type-specific effects for either CD4$^+$ T cells or monocytes, which were also significant in the main meQTL analysis. These associations involve 2,589 unique CpGs (1.1% of the CpGs with whole blood *cis*-meQTLs). At a more liberal threshold ($P \leq 0.05$), the number of CpGs influenced by genotype–cell type interactions increased slightly (3,916 CpGs, 1.6% of CpGs with cross-cell *cis*-meQTLs) (Supplementary table 2). Our results suggest that the majority of genetic effects that we detect on CpGs in whole blood are stable across different blood cell types. However, the frequency of a cell type in whole blood likely plays a role, where cell specific meQTLs for rare cell types may not to overlap whole blood meQTLs to the same extent, since DNA methylation levels are estimated in whole blood masking rarer cell-specific effects. The mean rations of CD4$^+$ T cells and monocytes across all samples were 0.199 ($sd = 0.073$) and 0.050 ($sd = 0.026$), respectively.

We observed that the cell type-specific meQTL effects also displayed a high heterogeneity across cohort sample. If we only consider results from the largest cohort sample (NCDS-99), the percentage of CpGs with whole blood *cis*-meQTLs slightly increases to 2.3% CpGs for CD4$^+$ T cells and 0.7% for monocytes (Supplementary table 2).

Our results overall show a smaller proportion of cell-specific meQTLs relative to whole blood meQTLs, and replicated $\approx 17\%$ of cell specific findings from previous work [10]. A relatively small number of CpGs had simultaneous cell-specific and whole blood meQTLs effects. From these observations, we conclude that the cell type-specific meQTL effects occurs at a lower extent than whole blood effects, and as expected tend to involve different

CpGs.

#### 1.7 Genetic regulation of methylation sites at enhancers

We performed an enrichment analysis to identify genomic annotations that are enriched or depleted for meQTLs associated with CpGs in enhancers. Overall, for most genetic annotations, the OR between meQTLs for CpGs within enhancers vs meQTLs for CpGs outside enhancers was very close to one for both *cis* and *trans* (although most of the *cis*-meQTLs were significant after Fisher's exact tests) (Supplementary fig. 11 and Supplementary table 9). That suggests that meQTLs for CpGs in enhancers have similar distribution patterns compared to all other meQTLs. As exceptions, we identified a considerable enrichment of *cis*-meQTLs in enhancers and TFBSs.

This result could be biased because *cis*-meQTL SNPs are more likely to be at enhancers due to the location of their associated CpGs. We repeated the enrichment analysis, taking as background set a thousand random samples of genetic variants, after categorizing all the SNPs according to their MAF and their distance from the EPIC CpGs only in enhancers (see Main methods). Interestingly, the distribution pattern on genetic annotations was almost identical to the enrichment analysis of meQTLs vs non-meQTL SNPs, presented in Figure 4 (Supplementary fig. 11 and Supplementary table 10). That confirmed that *cis*-meQTLs, either for CpGs in enhancers or not enhancers, are distributed equally across genetic annotations.

In the case of the previously mentioned enrichments in enhancers/TFBSs, we confirmed that this was not attributable to the location of the CpGs. In conclusion, CpGs in enhancers tend to be associated with meQTL SNPs in enhancers and TFBSs—more often than CpGs outside enhancers.

#### 1.8 Direction of effect in SMR associations between DNA methylation and gene expression

It is commonly accepted that methylation sites on promoters and enhancers are negatively correlated with the gene expression of the target gene, whereas CpG-sites within the gene bodies are positively correlated. In order to examine if our results from the SMR analysis with *cis*-meQTLs supported this assumption, we annotated the CpGs with regards to their position in the gene and compared the median of the $\beta_{\text{SMR}}$. We did the annotation with Ensembl version 104[9], accessed through the R package biomaRt[11].

In all regions, we observed median $\beta_{\text{SMR}}$ values below zero (Supplementary fig. 13). However, for the TSS200 sections (i.e. 200 bp upstream the transcription start site or TSS), which tend to be promoter regions, the negative shift was more pronounced. Altogether 64% of CpGs within TSS200 sites have a negative effect on the associated genes. Furthermore, intergenic regions showed a more symmetric bimodal distribution of their $\beta_{\mathrm{SMR}}$ values compared to the other genic positions.

#### 1.9 Integration of meQTL associations with Hi-C data

We investigated how many of the intra-chromosomal meQTL-CpG top pairs (*cis* and *trans*) were located in sites brought into proximity by three-dimensional (3D) conformations of the genome. For the annotation of the 3D genomic structures, we obtained data from Hi-C experiments published by Rao et al.[12] from the lymphoblastoid cell line GM12878 (primary experiment plus biological replicate, where available). The authors described three different types of 3D structures in which the genome is divided: a) contact domains, with enhanced contact frequency, of sizes ranging 40 kbp and 3 Mbp, b) six classes of larger subcompartments, which share distinctive histone marks, and c) chromatin loops, generally at the boundaries of contact domains.

We intersected the structural annotations with the meQTL associations to identify pairs where the CpG and the SNP were within the same contact domain or subcompartment. We also looked for associations where the SNP and the CpG were in the interacting regions through a DNA loop. For this, we focused on *cis* pairs with a minimum distance between the CpG and an SNP of 30 kbp—the minimum size of a loop—and intra-chromosomal *trans* pairs. For the overlap between the 3D annotations with the meQTL results, we used the R package GenomicRanges[13].

We found that 19.2% of CpGs with *cis*-meQTLs are within the same contact domain as the SNP and 32.7% in the same subcompartment. Only 0.04% are in interacting regions due to loop conformation. Regarding the intra-chromosomal *trans* associations, 0.2% of CpGs have a meQTL within the same contact domain and 9.1% in the same subcompartment. No CpG interacts through chromatin loops with *trans*-meQTLs.

We also looked for associations which may be in contact via topologically associated domains (TADs). We used TADs predicted by the 3D Genome Browser[14] from the same GM12878 Hi-C data[12]. 10.5% of CpGs with *cis*-meQTLs, and 3.9% of CpGs with intra-chromosomal *trans*-meQTLs are within the same TAD as the top SNP. Since the vast majority of intra-chromosomal *trans*-meQTLs are within 5 Mbp of the target CpG, they appear to be 'long-range' *cis*-meQTLs that TADs bring into physical proximity. To test this hypothesis, we compared the mean size of TADs with *cis* vs. *trans* associations. The mean size of the 2,636 TADs with CpGs with *cis*-meQTLs is 861.0 kbp, and of the 26 TADs with *trans* is 2.1 Mbp, which is a significative difference (two-tailed $t$-test, unequal variance assumed, $t_{(25.218)} = -6.4$, $P = 9.76 \times 10^{-7}$). Therefore, we conclude that the *trans*-meQTLs in TADs act as 'long-range' *cis*-meQTLs.

TADs predicted only from the GM12878 cell line are an incomplete proxy for whole blood domains, so we recalculated associations within TADs expanding the Hi-C source data to an alternate GM12878 experiment[15], plus spleen[16] and thymus[17], obtained from the 3D Genome Browser. The results remained consistent with the GM12878 data alone. Altogether, 17.1% (*cis*) and intra-chromosomal 36.5% (*trans*) of CpGs are annotated in the same TAD as their top meQTL. The size difference between TADs with *cis* and *trans* associations is also significant (mean TAD $\text{size}_{cis} = 1.2$ Mbp; mean TAD $\text{size}_{trans} = 3.4$ Mbp; two-tailed $t$-test, unequal variance assumed, $t_{(156.63)} = -8.0$, $P \leq 2.54 \times 10^{-13}$).

#### 1.10 Linkage disequilibrium (LD)-based clumping of meQTLs

We clumped meQTLs separately for *cis* and *trans* based on the LD between the SNPs. In the case of *cis* associations, we obtained 70,124 meQTL regions, with a mean of 65.7 SNPs per region ($sd = 126.3$). Each region covered in average 113.207 kbp ($sd = 168.461$). The number of *cis* associations of CpGs with unique clumped meQTL regions was 785,359.

For *trans*-meQTLs, LD clumping resulted in 3,108 regions with 77.5 meQTLs on average for each clump ($sd = 194.7$). The mean size of the meQTL regions was 123.688 kbp ($sd = 225.811$). There was a total of 8,976 unique associations of CpGs and *trans*-meQTL regions.

#### 1.11 MeQTL regulation in the major histocompatibility complex (MHC)

We analysed in detail the MHC region in chromosome 6 between the positions 28,477,797 and 33,448,354. This locus contains several highly regulated CpGs, such as cg13212186 with 47 *cis*-meQTLs and 7 *trans*-meQTLs. We quantified a modest enrichment in the MHC region for CpGs with *cis* associations (OR = 1.54, 95% CI [1.48–1.61]), and a high enrichment for CpGs with *trans* (OR = 9.37, 95% CI [8.55–10.25]). We observed that the MHC CpG-sites with meQTLs have contrasting enrichment/depletion patterns compared to the whole set of CpGs with meQTLs (Supplementary fig.16a and Supplementary table 14). For example, promoters in the MHC frequently contain CpGs with *cis*-meQTLs, opposed to promoters from the rest of the genome. The genetic variants associated with the MHC CpGs also has slightly different patterns compared to all others meQTLs. The largest difference is that gene bodies are depleted, and intergenic regions are enriched for meQTLs of MHC CpGs, in contrast to patterns observed for genome-wide meQTLs (Supplementary fig. 16b and Supplementary table 15).

For both *cis* and *trans* associations, the key regulatory regions with most associations are located in the MHC locus, with as many as 1,949 and 360 associations respectively. We detected an extensive overrepresentation of meQTL SNPs in the MHC for both *cis* (OR = 12.93, 95% CI [12.16–13.77]) and *trans* (OR = 28.16, 95% CI [27.56–28.73]), compared to all the available SNPs for the analysis. CpGs that were in *trans*-associations with MHC meQTLs tend to be more frequently in genic and coding region—compared to CpGs with meQTLs outside the MHC (Supplementary fig. 17 and Supplementary table 16).

#### 1.12 Sensitivity analyses

Recently, the Genetics of DNA Methylation Consortium (GoDMC) published the largest meQTL analysis to date in the 450K array, comprising 27,750 samples[18]—more than ten times the sample of our study. This, together with the doubling of the methylome coverage in the EPIC array, limited statistical power of our study to detect meQTLs compared to GoDMC. To overcome this drawback, we split the EPIC CpGs into two sets for meQTL sensitivity analyses: 1) legacy 450K probes only for replication of GoDMC results, and 2) novel EPIC-only probes for detection of new associations. We performed genome-wide association analysis, meta-analysis and permutations to determine signals with a FDR $\leq 0.05$ in each set separately. The estimated thresholds of $P$-values were almost equal for the split analysis compared to the main analysis (for the 450K legacy probes set $P_{cis} \leq 2.23 \times 10^{-4}$ and $P_{trans} \leq 4.87 \times 10^{-9}$; for the novel EPIC-only probes set $P_{cis} \leq 2.18 \times 10^{-4}$ and $P_{trans} \leq 1.84 \times 10^{-9}$). As a result, the difference in the number of significant associations with the main analysis is negligible.

To confirm the validity of the multiple comparisons of means based on Student's $t$-test, we performed a Mann-Whitney $U$ tests as sensitivity analyses. This, as we were aware that some of the assumptions of the $t$-test might not be fully met by the datasets. The only assumption of the Mann-Whitney test is the independence of the datasets compared. In all cases, the results of the Mann-Whitney test were consistent with those of the respective $t$-test (Supplementary table 4).

### 2 Cohort-specific acknowledgements

#### 2.1 TwinsUK

The TwinsUK study was funded by the Welcome Trust; European Community's Seventh Framework Programme (FP7/2007-2013). The study also receives support from the National Institute for Health Research (NIHR)-funded BioResource, Clinical Research Facility and Biomedical Research Centre based at Guy's and St Thomas' NHS Foundation Trust in partnership with King's College London. The DNA methylation TwinsUK data and analysis included in this study was funded in part from the UK Economic and Social Research Council (ESRC ES/N000404/1 to JTB) and JPI-HDHL DIMENSION project, funded in the UK via the Biotechnology and Biological Sciences Research Council (BBSRC BB/S020845/1 and BB/T019980/1 to JTB).

#### 2.2 The MRC National Survey of Health and Development or 1946 British birth cohort

The U.K. Medical Research Council provides core funding for the MRC National Survey of Health and Development (MC_UU_00019/1). KKO is supported by the Medical Research Council (MC_UU_00006/2). We acknowledge study members for their lifelong participation and past and present members of the study teams, including members of the MRC Epidemiology unit in Cambridge, who helped to collect and process the data.

#### 2.3 The National Child Development Study or 1958 British birth cohort

We acknowledge the co-operation and participation of the individuals who voluntarily participate in the 1958 birth cohort study. We thank the Economic and Social Research Council for funding these cohorts through the Centre for Longitudinal Studies (CLS) at the UCL Institute of Education, London. We thank the Economic and Social Research Council for funding the Cross Cohort Research Programme (CCRP) (grant number: ES/M008584/1). Blood collection for the 1958 cohort was funded by the Medical Research Council (MRC) (grant G0000934 to the clinical examination and DNA banking of the 1958 cohort). CR acknowledges funding for the 1958BC-2 DNA methylation from ESRC ES/N000498/1. We like to thank a large number of stakeholders from academic, policy-maker and funder communities and colleagues at CLS involved in data collection and management.

### 3 Supplementary tables

**Supplementary table 1. Number of unique genotype variants, DNA methylation probes and pairs analysed and identified as candidates per cohort dataset.**

| Cohort | Input[a] | | *Cis* candidates[b] | | | *Trans* candidates[c] | | |
|---|---|---|---|---|---|---|---|---|
| | SNPs | CpGs | SNPs | CpGs | Pairs | SNPs | CpGs | Pairs |
| TwinsUK | 5,598,605 | 723,814 | 4,998,690 | 560,241 | 47,008,415 | 3,663,233 | 498,097 | 8,558,062 |
| 1946BC-99 | 5,768,798 | 713,766 | 5,613,022 | 631,388 | 123,047,963 | 5,251,132 | 704,712 | 30,666,785 |
| 1946BC-09 | 5,696,058 | 713,766 | 5,076,588 | 593,338 | 43,498,983 | 4,722,131 | 702,725 | 21,148,528 |
| 1958BC-1 | 5,800,303 | 723,675 | 5,238,058 | 609,327 | 47,294,158 | 5,283,752 | 716,385 | 22,000,014 |
| 1958BC-2 | 5,811,581 | 699,968 | 4,516,795 | 559,841 | 17,128,256 | 5,160,748 | 693,353 | 20,611,728 |
| All | 6,361,063 | 724,499 | 6,283,304 | 720,128 | 189,202,234 | 6,314,322 | 724,460 | 100,814,822 |

[a] Number of unique SNPs/CpGs that were analysed in each cohort, after quality control filtering.
[b] Candidate associations for following meta-analysis at a $P_{cis} \leq 2.21 \times 10^{-4}$.
[c] Candidate associations for following meta-analysis at a $P_{trans} \leq 3.35 \times 10^{-9}$.

**Supplementary table 2. Cell type-specific meQTLs for $CD4^+$ T cells and monocytes.**

| Cell type | Results from | $P \leq 0.05$ | | | $P \leq 2.21 \times 10^{-4}$ | | |
|---|---|---|---|---|---|---|---|
| | | **SNPs** | **CpGs** | **Pairs** | **SNPs** | **CpGs** | **Pairs** |
| **Considering only overlapping associations with whole blood** | | | | | | | |
| $CD4^+$ T | Meta-analysis | 79,398 | 3,000 | 94,144 | 56,217 | 2,015 | 65,401 |
| Monocytes | Meta-analysis | 10,107 | 1,098 | 10,316 | 5,318 | 637 | 5,358 |
| $CD4^+$ T + Monocytes | Meta-analysis | 87,208 | 3,916 | 102,671 | 60,763 | 2,589 | 70,321 |
| $CD4^+$ T | NCDS-99 samples[a] | | | | 178,819 | 5,711 | 232,080 |
| Monocytes | NCDS-99 samples[a] | | | | 27,698 | 1,803 | 30,226 |
| **Total associations** | | | | | | | |
| $CD4^+$ T | Meta-analysis | 300,826 | 40,663 | 335,781 | 199,439 | 26,990 | 217,380 |
| Monocytes | Meta-analysis | 351,897 | 57,280 | 385,291 | 237,193 | 39,697 | 251,602 |
| $CD4^+$ T + Monocytes | Meta-analysis | 616,821 | 93,716 | 718,920 | 699,031 | 64,692 | 468,372 |
| $CD4^+$ T | NCDS-99 samples[a] | | | | 178,819 | 75,422 | 879,842 |
| Monocytes | NCDS-99 samples[a] | | | | 585,147 | 75,488 | 685,491 |

[a] Results at $P \leq 0.05$ are not reported for NCDS-99, since in the meQTL analysis for each independent sample cohort only associations that reached $P \leq 5 \times 10^{-3}$ were stored.

**Supplementary table 3. MeQTL replication results for selected CpG sites using MeDIP-seq.**

| CpG | Notes | Original meQTLs | | Replicated meQTLs[a] | | Replication rate (%)[b] | |
|---|---|---|---|---|---|---|---|
| | | ***Cis*** | ***Trans*** | ***Cis*** | ***Trans*** | ***Cis*** | ***Trans*** |
| cg25014118 | Highly regulated CpG ($n_{cis} = 50$, $n_{trans} = 6$) | 4,712 | 1,816 | 32 | 0 | 1 | 0 |
| cg00128506 | Highly regulated CpG ($n_{cis} = 48$, $n_{trans} = 13$) | 4,550 | 1,979 | 3,696 | 1,605 | 81 | 81 |
| cg18111489 | Highly regulated CpG ($n_{cis} = 47$, $n_{trans} = 3$) | 4,429 | 1,762 | 0 | 0 | 0 | 0 |
| cg16423305 | Highly regulated CpG ($n_{cis} = 42$, $n_{trans} = 21$) | 3,889 | 2,508 | 1,635 | 1,205 | 42 | 48 |
| cg07143125 | CpG with strongest *cis*-meQTL association (rs61914738, $P \to 0$, $\beta = 2.73$) | 1,254 | 4,184 | 0 | 0 | 0 | 0 |
| cg13904258 | CpG with strongest *cis*-meQTL association (rs35441453, $P \to 0$, $\beta = 2.34$) | 307 | 0 | 256 | 0 | 83 | – |
| cg00918944 | CpG with strongest *cis*-meQTL association (rs2236325, $P \to 0$, $\beta = 2.28$) | 87 | 0 | 45 | 0 | 52 | – |
| cg05808124 | CpG with strongest *trans*-meQTL association (rs57812432, $P \to 0$, $\beta = -1.59$) | 2,44 | 94 | 238 | 0 | 98 | 0 |
| cg11024963 | CpG with more QTL co-localisations with gene expression ($n_{\text{SMR}} = 13$) | 40 | 0 | 19 | 0 | 48 | – |
| cg06162668 | CpG with more QTL co-localisations with phenotypes ($n_{\text{SMR}} = 6$) | 367 | 0 | 236 | 0 | 64 | – |
| Total | | 19,879 | 12,343 | 6,157 | 2,810 | 31 | 23 |

[a] At a $P < 0.005$ and after combining the results from MeDIP-seq bins covering a single site.
[b] Number of replicated meQTLs divided by the number of original meQTLs.

**Supplementary table 4. Mann-Whitney *U* test as sensitivity analysis for *t*-test.**

| Data | *U* statistic | *P*-value[a] | Difference in location[b] [95% CI] |
|---|---|---|---|
| Heritabilities of EPIC-only vs. 450K legacy probes | $6.72 \times 10^{10}$ | $< 2.2 \times 10^{-16}$ | $2.59 \times 10^{-5}$ $[5.97 \times 10^{-6}, 5.94 \times 10^{-5}]$ |
| Proportion of variance explained by *cis* vs. *trans* meQTL effects | $7.8 \times 10^{12}$ | $< 2.2 \times 10^{-16}$ | $-0.042$ $[-0.0423, -0.0421]$ |
| Proportion of variance explained by meQTLs in CpGs with *cis*+*trans* effects vs. *cis*/*trans* only | $3.74 \times 10^{8}$ | $< 2.2 \times 10^{-16}$ | $0.094$ $[0.090, 0.099]$ |
| Heritabilities of CpGs with no meQTL effects vs. CpGs with meQTL effects. | $3.48 \times 10^{10}$ | $< 2.2 \times 10^{-16}$ | $-0.112$ $[-0.113, -0.111]$ |

[a] Normal approximation
[b] Estimate of the median of the difference between a sample from $x$ and a sample from $y$

### 4 Supplementary figures

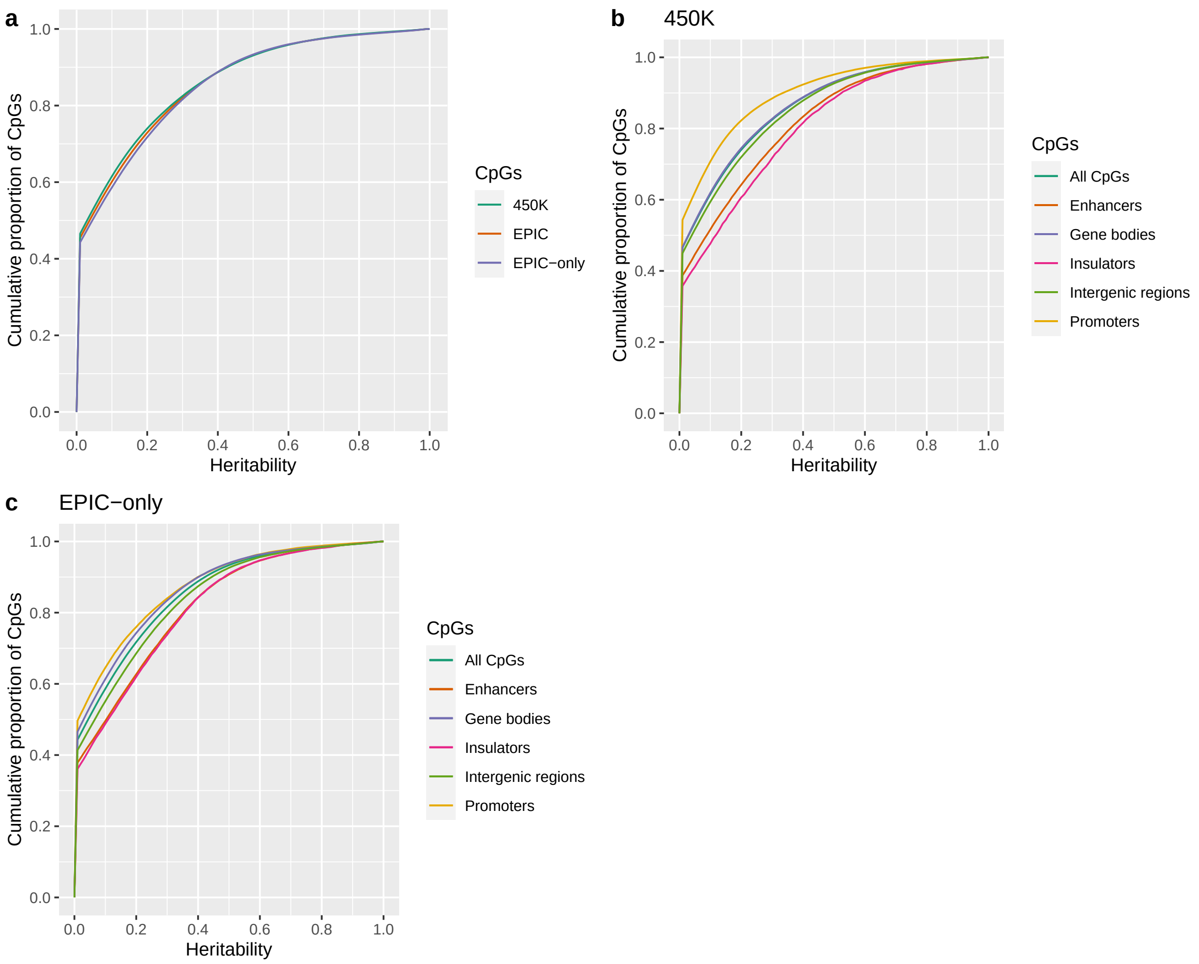

**Supplementary figure 1. Cumulative proportion of heritability estimates by probe type from the EPIC array.** (**a**) Heritability results for 450K array legacy probes (375,336 probes) are shown in green, for the novel EPIC-only probes (348,091 probes) in purple, and for the full set of EPIC array probes (723,814 probes) in orange. (**b**) Cumulative proportion of heritability estimates by genomic annotations for 450K array legacy probes. (**c**) Cumulative proportion of heritability estimates by genomic annotations for the novel EPIC-only probes.

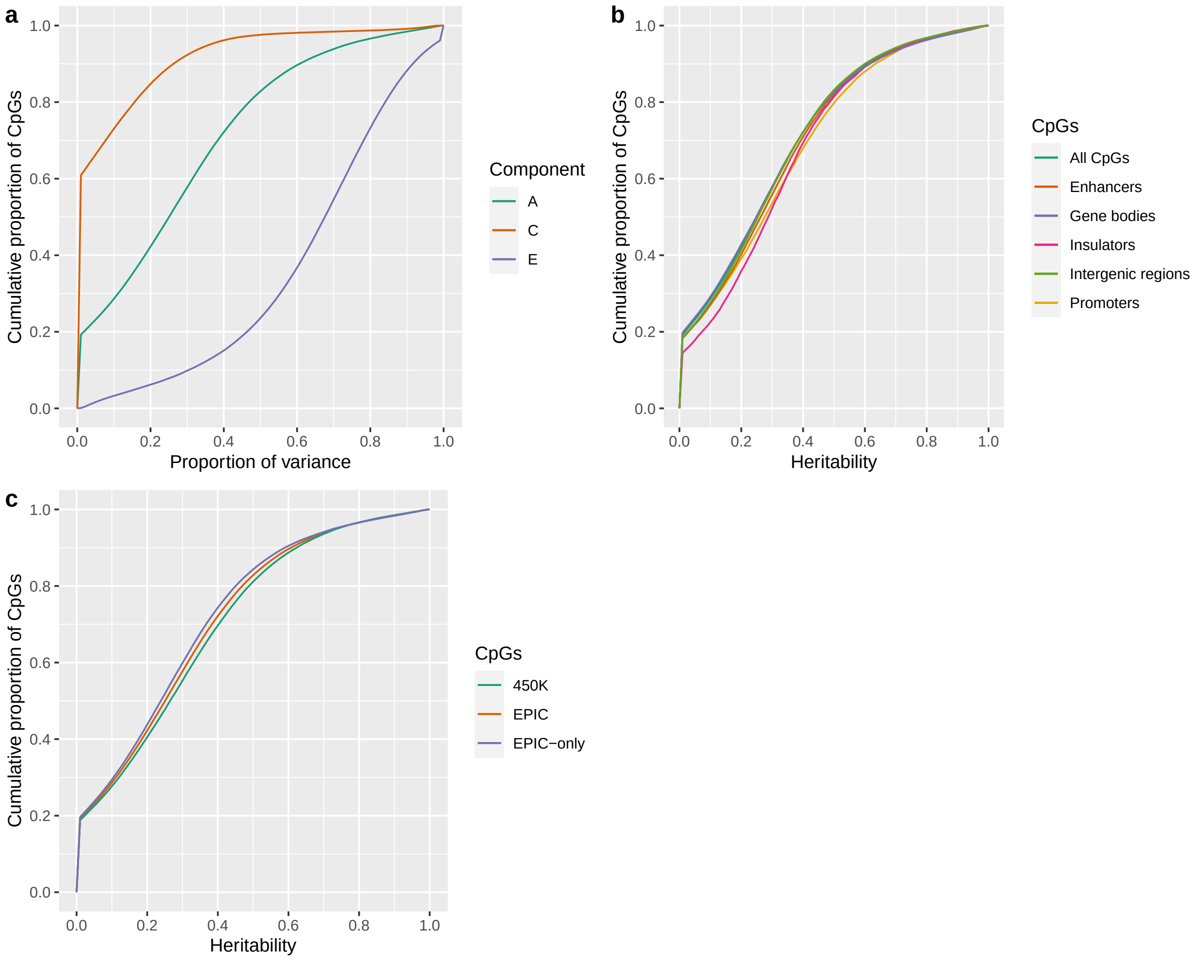

**Supplementary figure 2. Proportion of variance of DNA methylation levels attributed to genetic variation for variable CpG sites.** Estimates for the 238,988 CpGs sites with methylation $\beta$-values $sd > 0.025$, after a classical twin study of 88 MZ and 70 DZ twin pairs from the TwinsUK cohort. (**a**) Cumulative proportion of variance components of the ACE model. (**b**) Cumulative proportion of heritability estimates by genomic annotations. (**c**) Cumulative proportion of heritability estimates by probe type from the EPIC array.

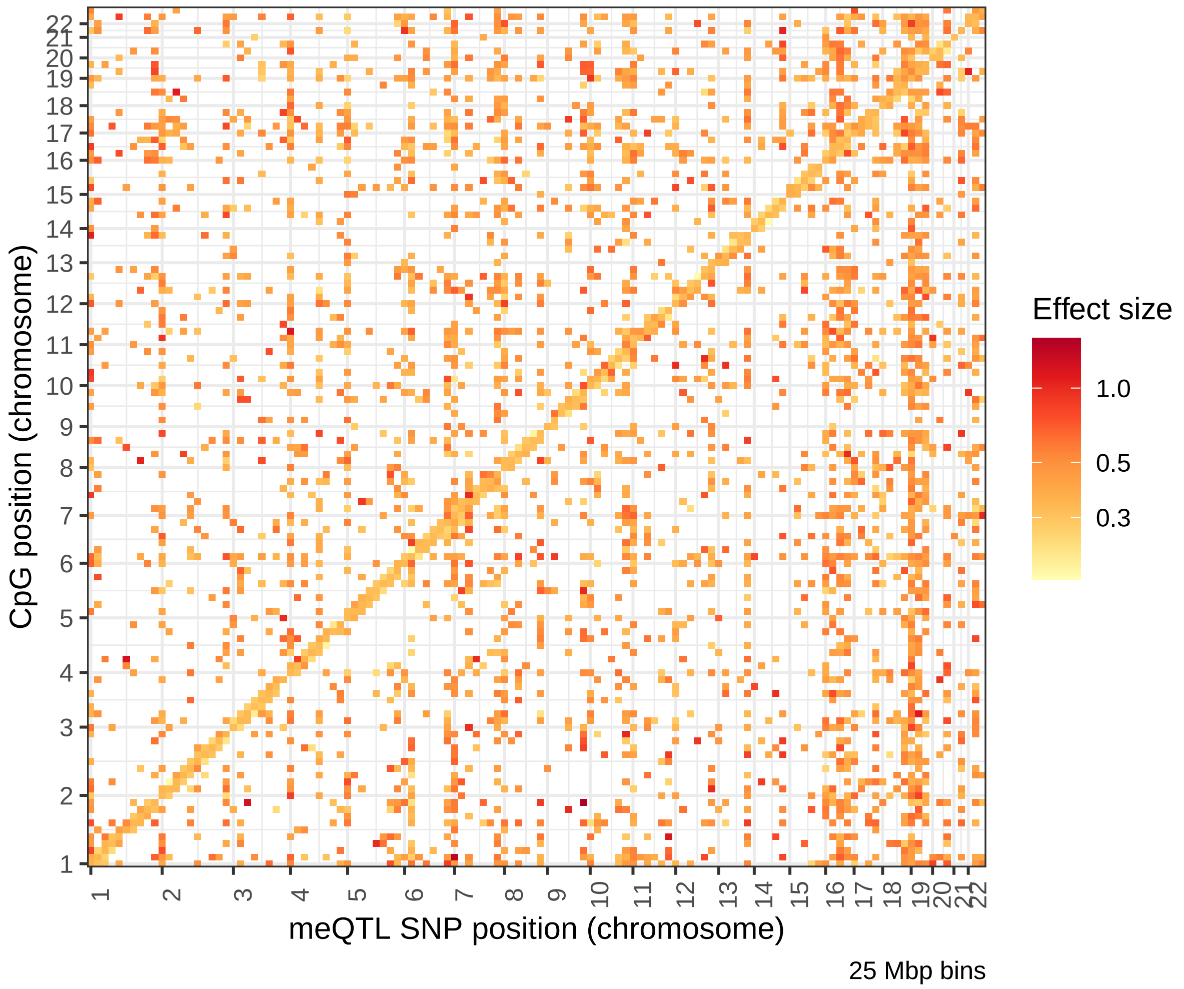

**Supplementary figure 3. Absolute effect sizes of meQTLs for CpG sites genome-wide.** MeQTL associations at a significance level of FDR $\leq 0.05$. The $x$-axis corresponds to the position of the SNPs within in the 22 chromosomes and the $y$-axis to the position of the CpGs, with each pixel binning a range of 25 Mbps. The colour scale indicates the inverse variance-weighted average of the absolute effect sizes of the associations between those specific CpGs/SNPs locations, on logarithmic scale.

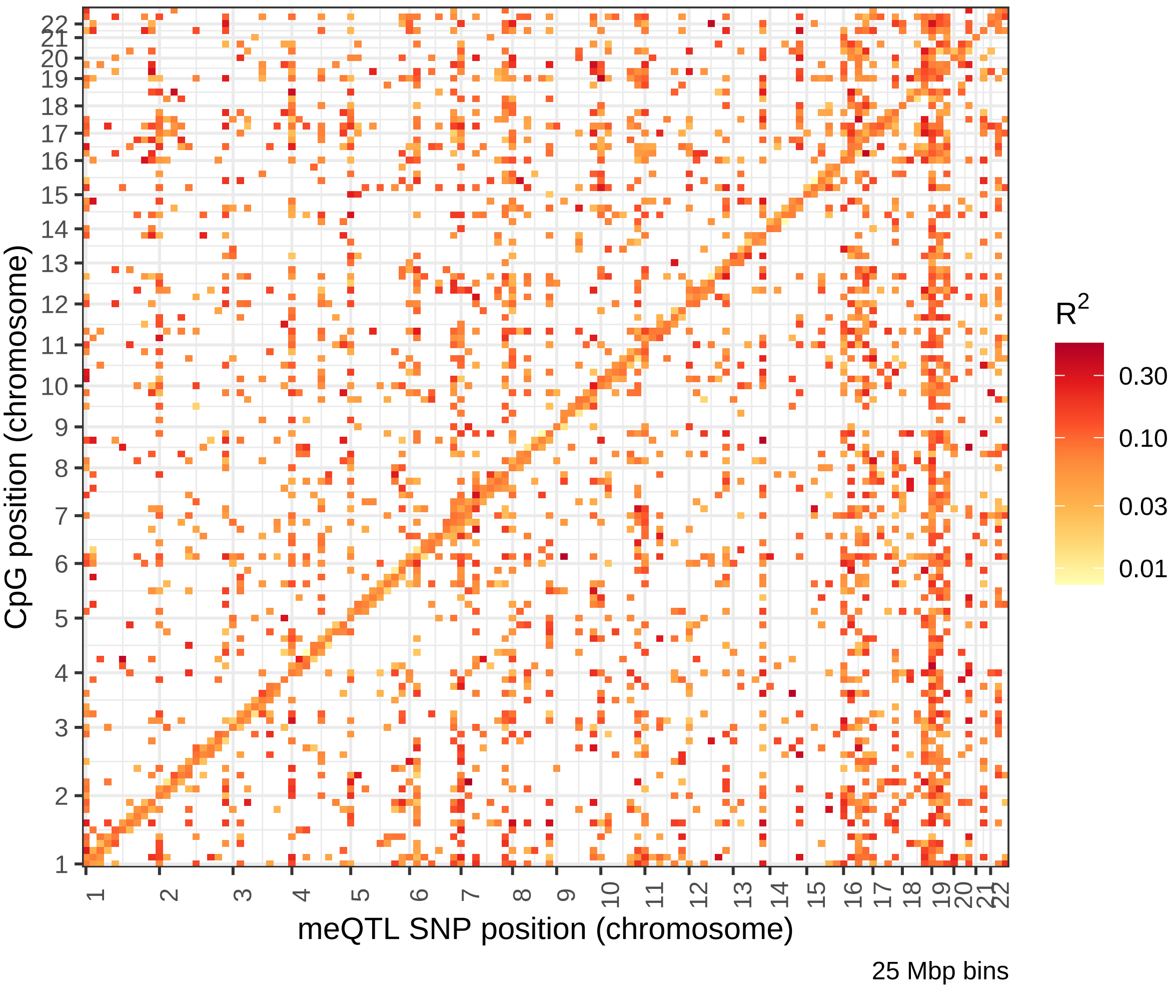

**Supplementary figure 4. Explained variance of meQTLs for CpG sites genome-wide.** MeQTL associations at a significance level of FDR $\leq 0.05$. The $x$-axis corresponds to the position of the SNPs within in the 22 chromosomes and the $y$-axis to the position of the CpGs, with each pixel binning a range of 25 Mbps. The colour scale indicates the average $R^2$ of the associations between those specific CpGs/SNPs locations, on logarithmic scale. $R^2$ of meQTL associations was estimated in the candidate associations from the 1946BC-99 sample.

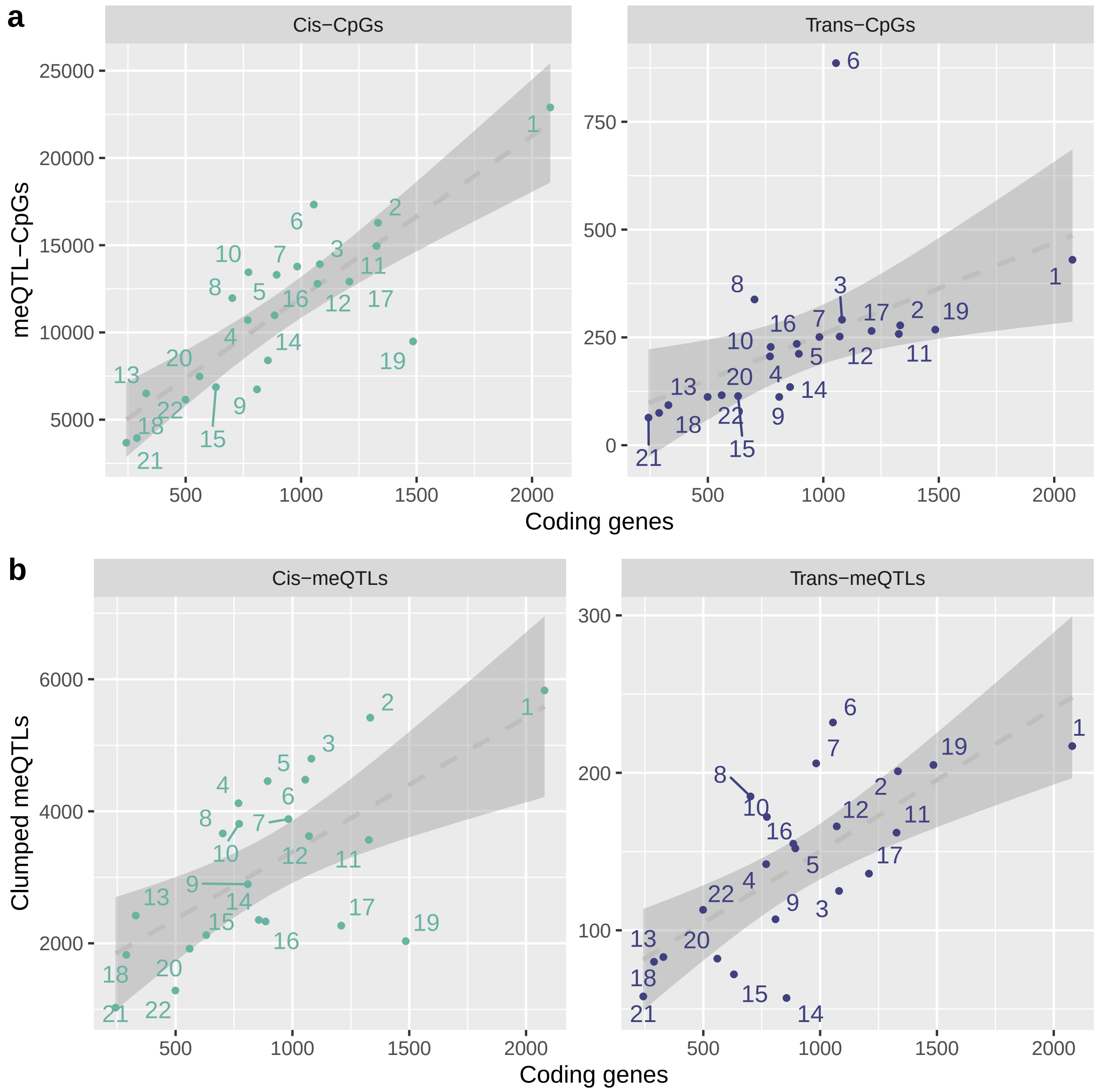

**Supplementary figure 5. Chromosomal distribution of CpGs and meQTL SNPs.** Dots represent values for each chromosome. On the $x$-axis, the number of coding genes (according to Ensembl version 104). On the $y$-axis, the number of (**a**) CpGs with meQTL associations(significance level of FDR $\leq 0.05$), or (**b**) LD-clumped meQTL regions. The dashed line shows the linear regression between the two variables and the shaded region, the 95% confidence bands.

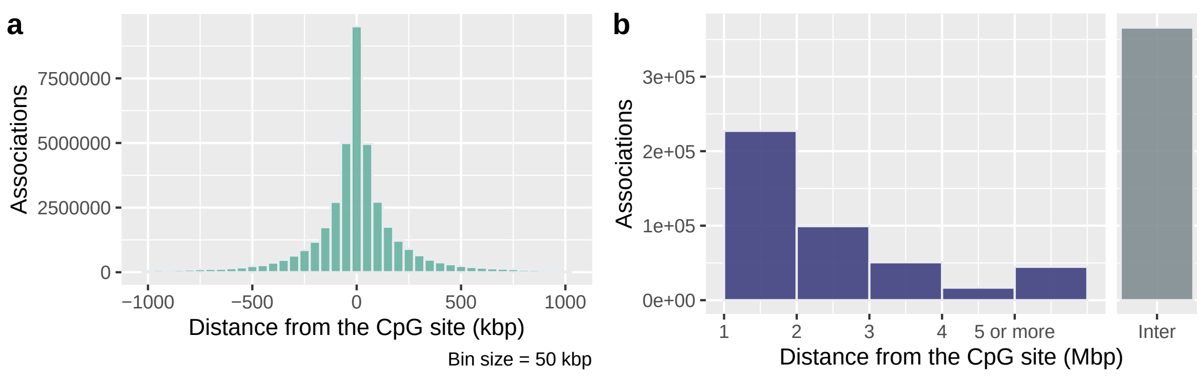

**Supplementary figure 6. Distances between the CpGs and their meQTL SNPs.** (**a**) Histogram of distances, considering the full set of associations. (**b**) Bar plot of absolute distances, considering the full set of associations. Intra-chromosomal associations are shown in purple, and inter-chromosomal in grey.

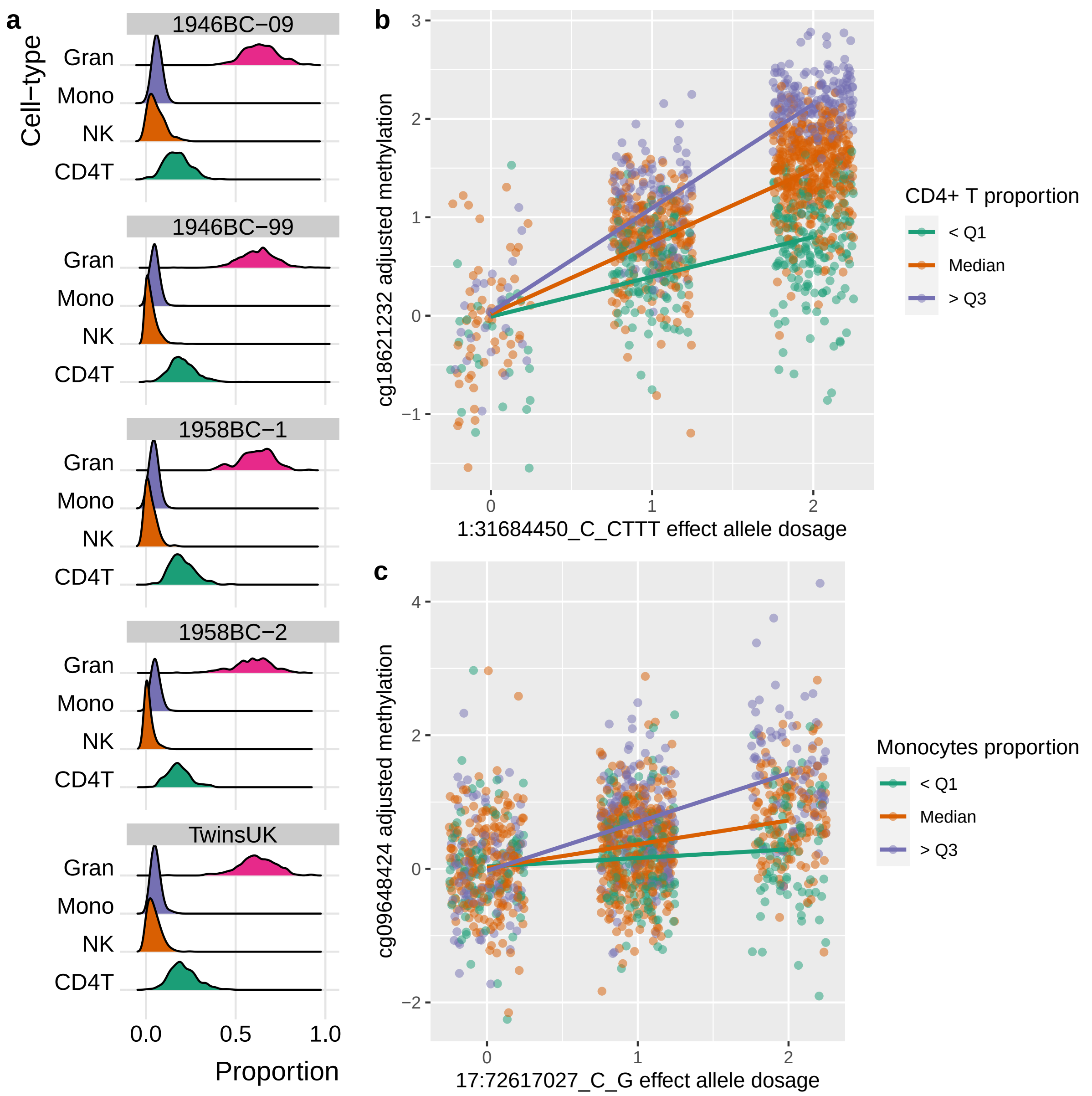

**Supplementary figure 7. Cell type-specific meQTLs analyses**. (**a**) Distribution of cell composition in whole blood samples by cohort. Cell types used as covariates in the main meQTL analysis and estimated with the method described by Houseman et al.[19] are shown (Gran = granulocytes, Mono = monocytes, NK = natural killer, CD4T = $CD4^+$ T cells). (**b**) Most significant interaction between a meQTL and $CD4^+$ T cell levels, or (**c**) monocytes. The $x$-axes show the dose of the effect allele of the meQTL SNP, the $y$-axes show the DNA methylation levels CpG site adjusted for all covariates (including the main meQTL effect and the main cell effect), the colours show cell proportion discretised by quartiles (for visualisation purposes only), and the lines show the linear regressions between the two variables in each cell proportion group. For visualisation purposes, only samples from the NCDS-99 cohort are shown.

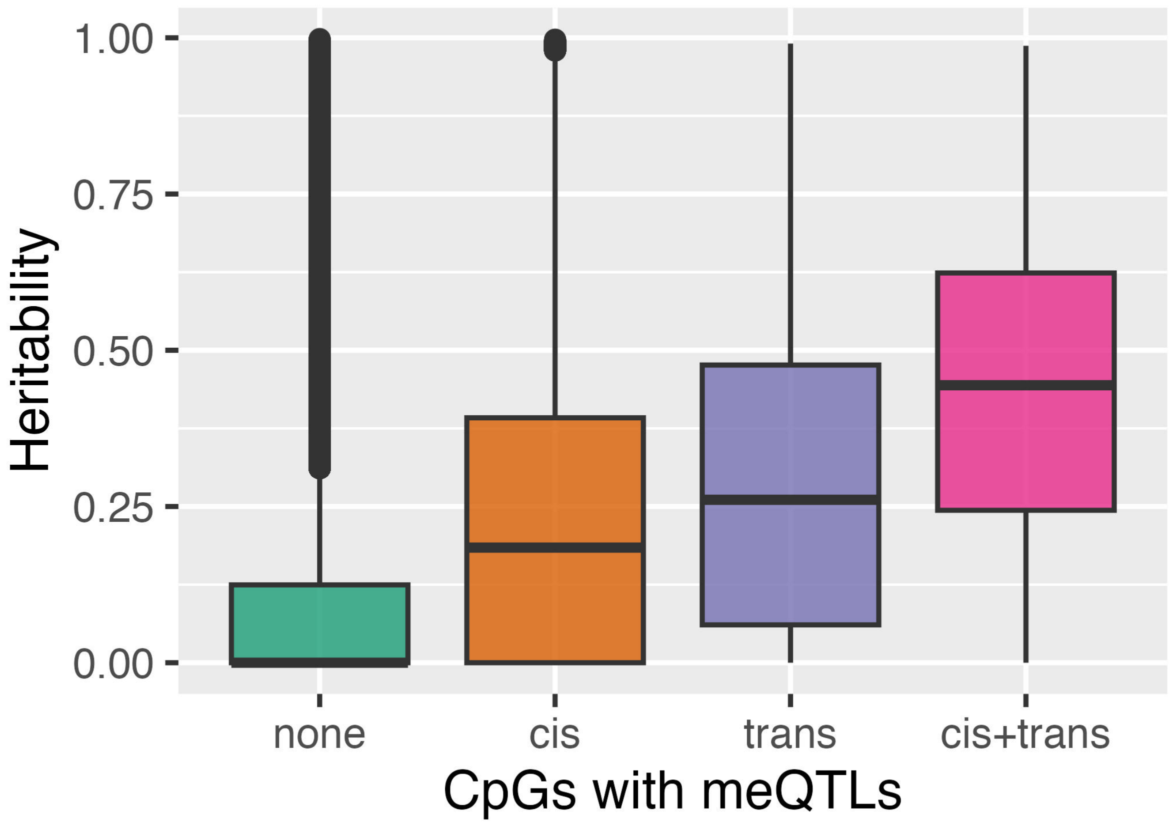

**Supplementary figure 8. Heritability of CpGs with meQTLs.** CpGs are grouped for the boxplot depending on whether they have *cis*-meQTLs, *trans*, or simultaneously *cis* and *trans* associations (significance level of FDR $\leq 0.05$).

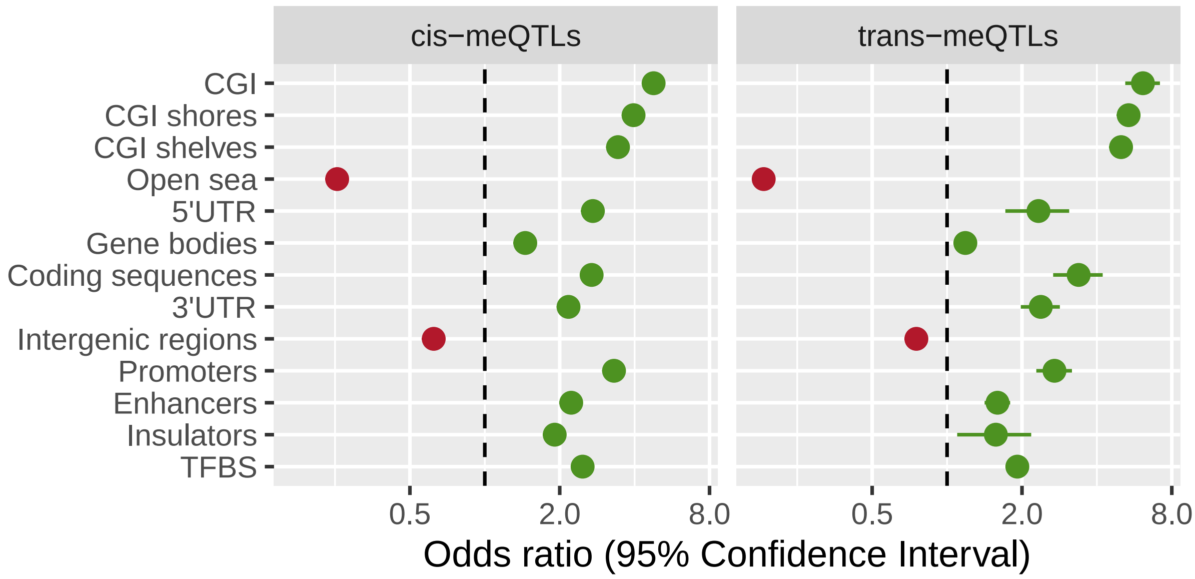

**Supplementary figure 9. Enrichment in genomic annotations of meQTL SNPs.** The $x$-axis indicates the odds ratio and its 95% confidence interval (in logarithmic scale) for meQTL SNPs located within a specific genomic annotation. Background SNPs for comparison were the full panel of genetic variants. Significant enrichment is marked in green, depletion in red, and non-significant genomic annotations in grey.

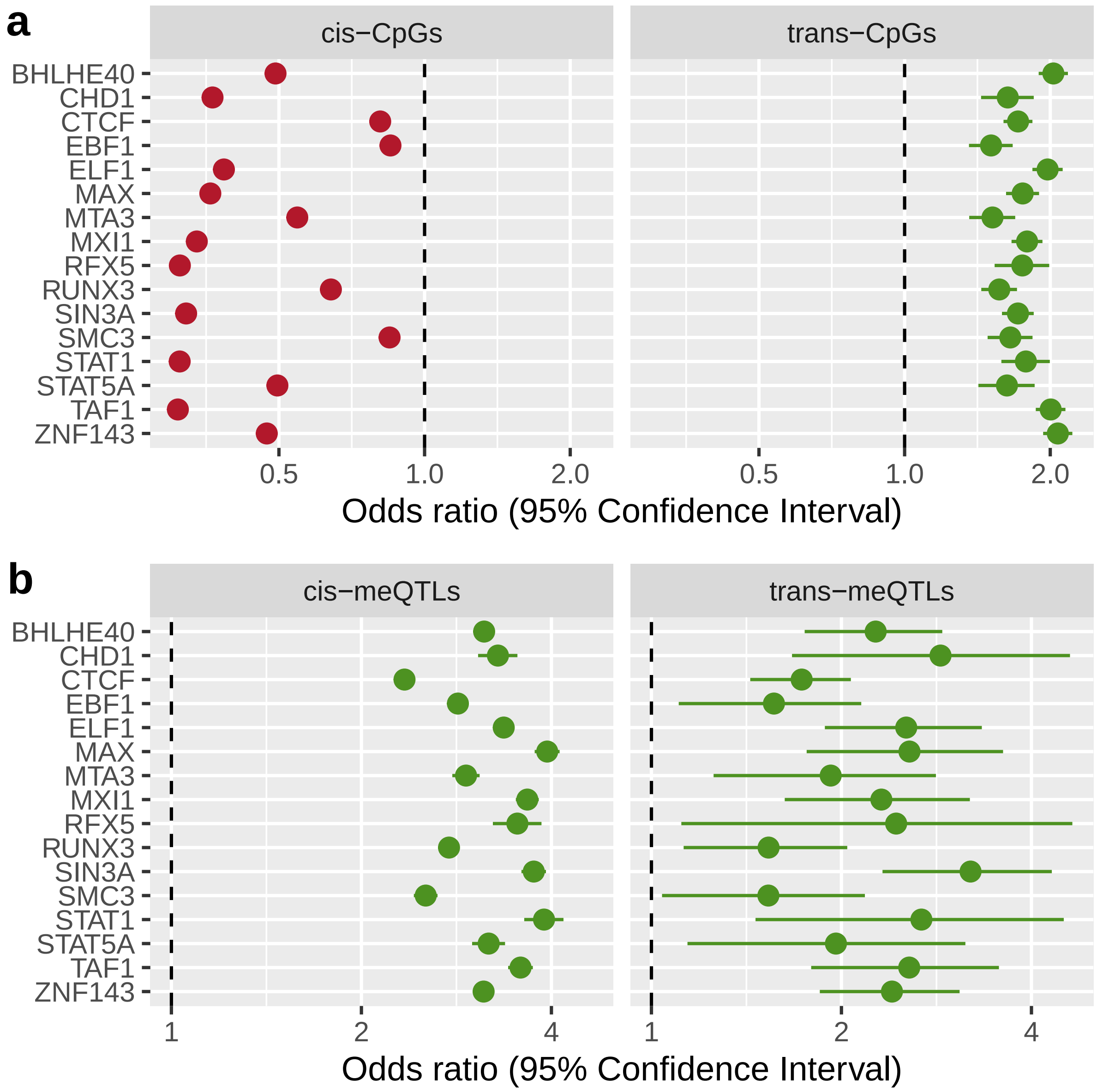

**Supplementary figure 10. Enrichment in TFBSs of meQTL SNPs and their CpGs.** The $x$-axis indicates the odds ratio and its 95% confidence interval (in logarithmic scale) for (**a**) CpGs with meQTLs, or (**b**) meQTL SNPs, located within a specific binding site of selected transcription factors. Significant enrichment is marked in green, depletion in red, and non-significant genomic annotations in grey.

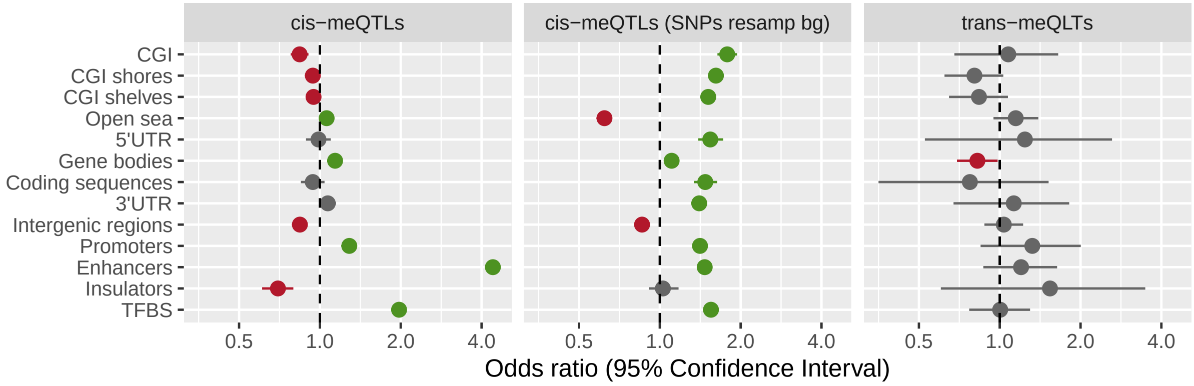

**Supplementary figure 11. Enrichment in genomic annotations of meQTL SNPs associated with CpGs in enhancers.** The $x$-axis indicates the odds ratio and its 95% confidence interval (in logarithmic scale) for meQTL SNPs located within a specific genomic annotation. Background SNPs for comparison were the full set of meQTLs with the lowest $P$ (side panels), or a random sample (middle panel). Significant enrichment is marked in green, depletion in red, and non-significant genomic annotations in grey.

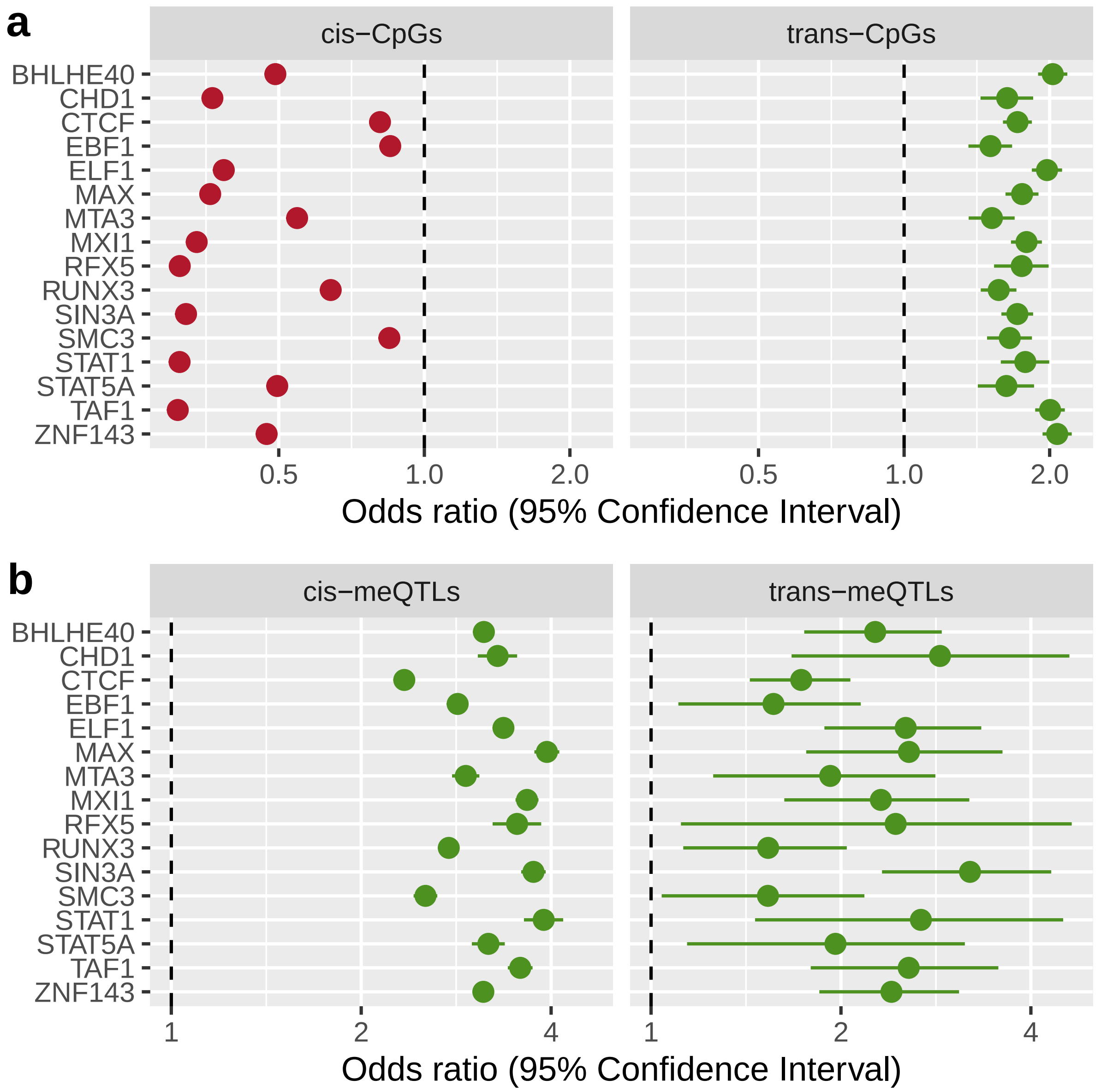

**Supplementary figure 12. Enrichment in genomic annotations of CpGs associated with nearby genes after SMR.** The $x$-axis indicates the odds ratio and its 95% confidence interval (in logarithmic scale) for CpGs with *cis*-meQTL co-localised with *cis*-eQTLs, located within a specific genomic annotation. Background CpGs for comparison were the full set of CpGs with meQTLs. Significant enrichment is marked in green, depletion in red, and non-significant genomic annotations in grey.

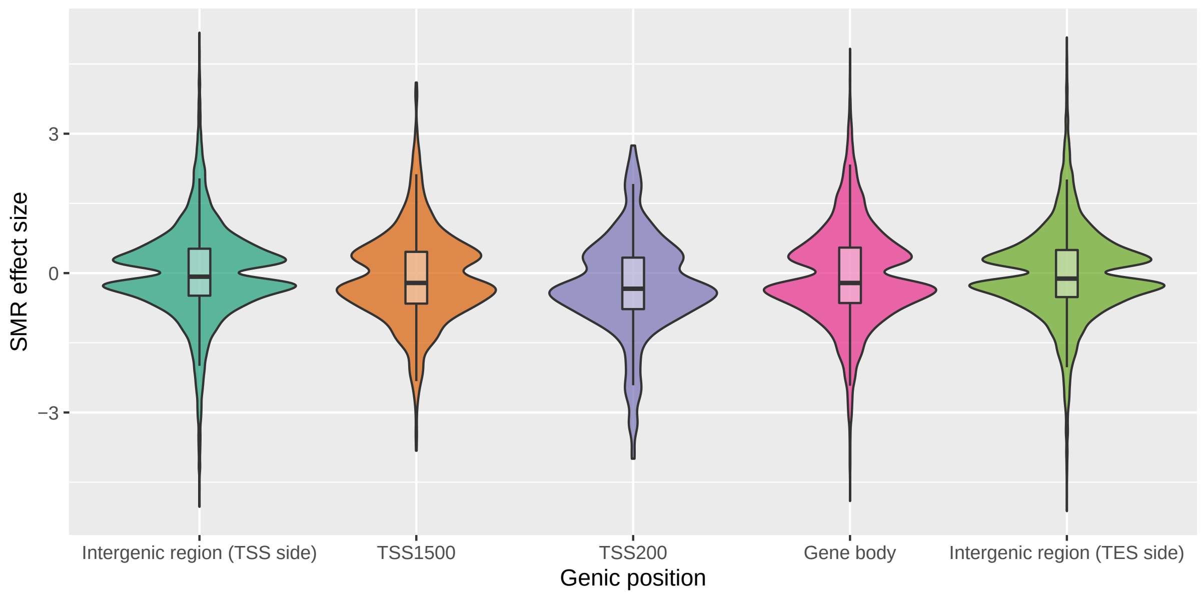

**Supplementary figure 13. Estimated SMR effect sizes of CpGs associated with nearby genes through co-localised QTLs.** Associations are grouped for the violin plot depending on the location of the CpG with respect to the associated gene. TSS1500/TSS200 refers to the 1500/200 bp regions upstream of the transcription start sites.

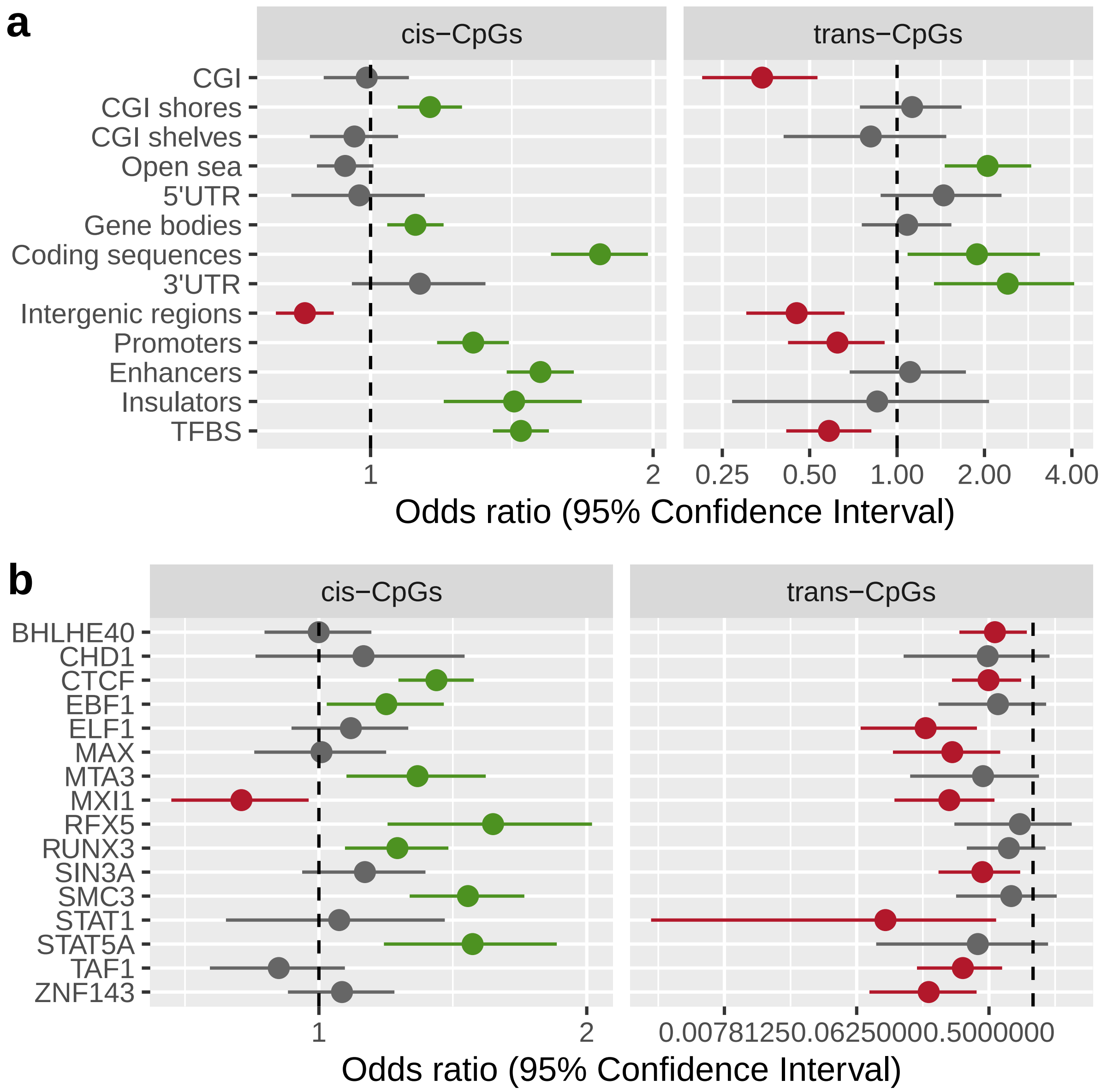

**Supplementary figure 14. Enrichment in genomic annotations and TFBSs of highly regulated CpGs.** The $x$-axis indicates the odds ratio and its 95% confidence interval (in logarithmic scale) for highly regulated CpGs with meQTLs, located within a specific (**a**) genomic annotation, or (**b**) binding site of selected transcription factors. Background CpGs for comparison were the full set of CpGs with meQTLs. Significant enrichment is marked in green, depletion in red, and non-significant genomic annotations in grey.

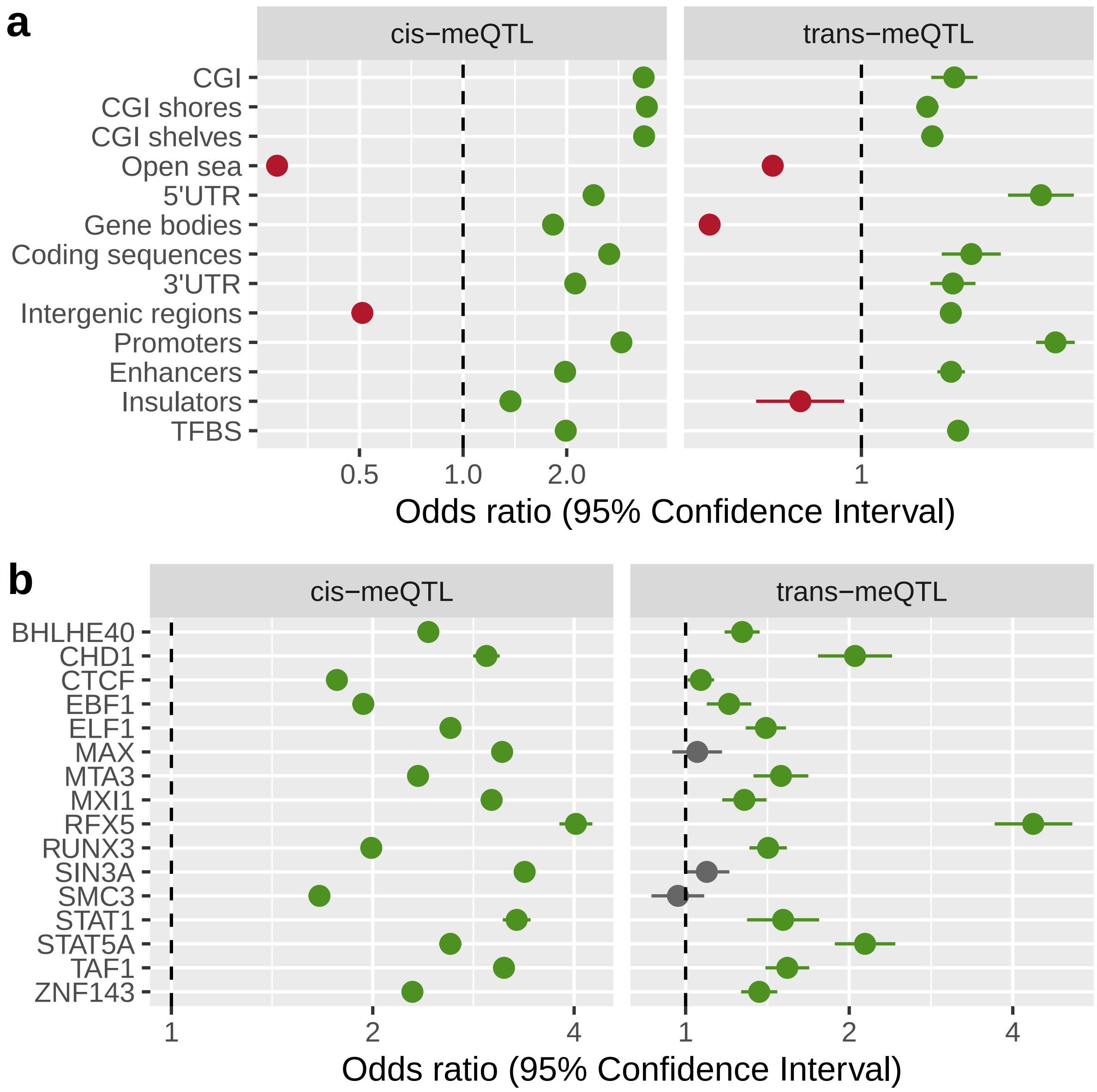

**Supplementary figure 15. Enrichment in genomic annotations and TFBSs of SNPs in key regulatory meQTL regions.** The $x$-axis indicates the odds ratio and its 95% confidence interval (in logarithmic scale) for SNPs in highly regulatory regions, located within a specific (**a**) genomic annotation, or (**b**) binding site of selected transcription factors. Background SNPs for comparison were the full set of meQTLs. Significant enrichment is marked in green, depletion in red, and non-significant genomic annotations in grey.

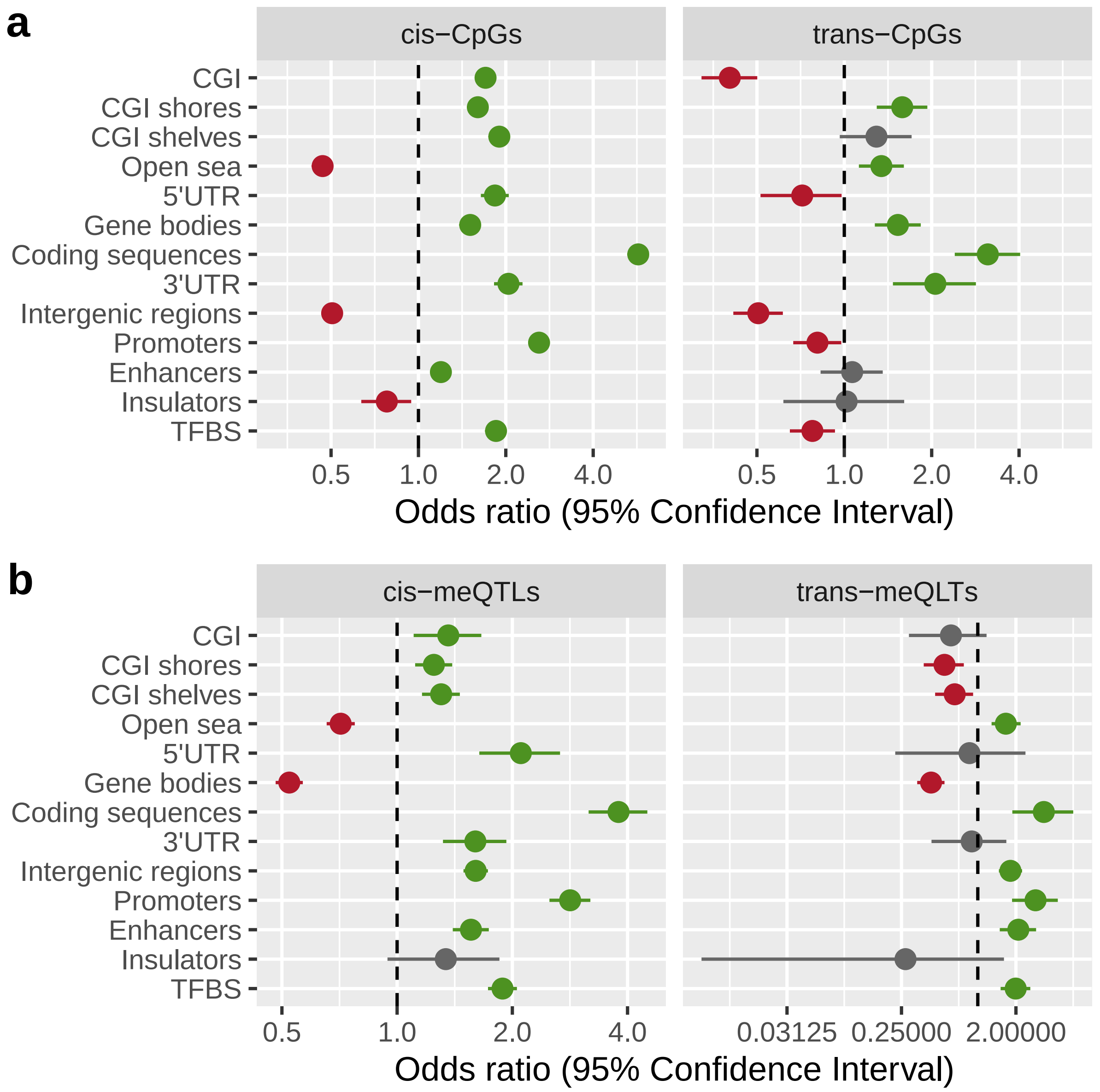

**Supplementary figure 16. Enrichment in genomic annotations of CpGs in the MHC region and their meQTL SNPs.** The $x$-axis indicates the odds ratio and its 95% confidence interval (in logarithmic scale) for (**a**) CpGs in the MHC region with meQTLs, or (**b**) meQTL SNPs of CpGs in the MHC region, located within a specific genomic annotation. Background CpGs for comparison were the full set of CpGs with meQTLs. Background SNPs for comparison were the full set of meQTLs with the lowest *P*. Significant enrichment is marked in green, depletion in red, and non-significant genomic annotations in grey.

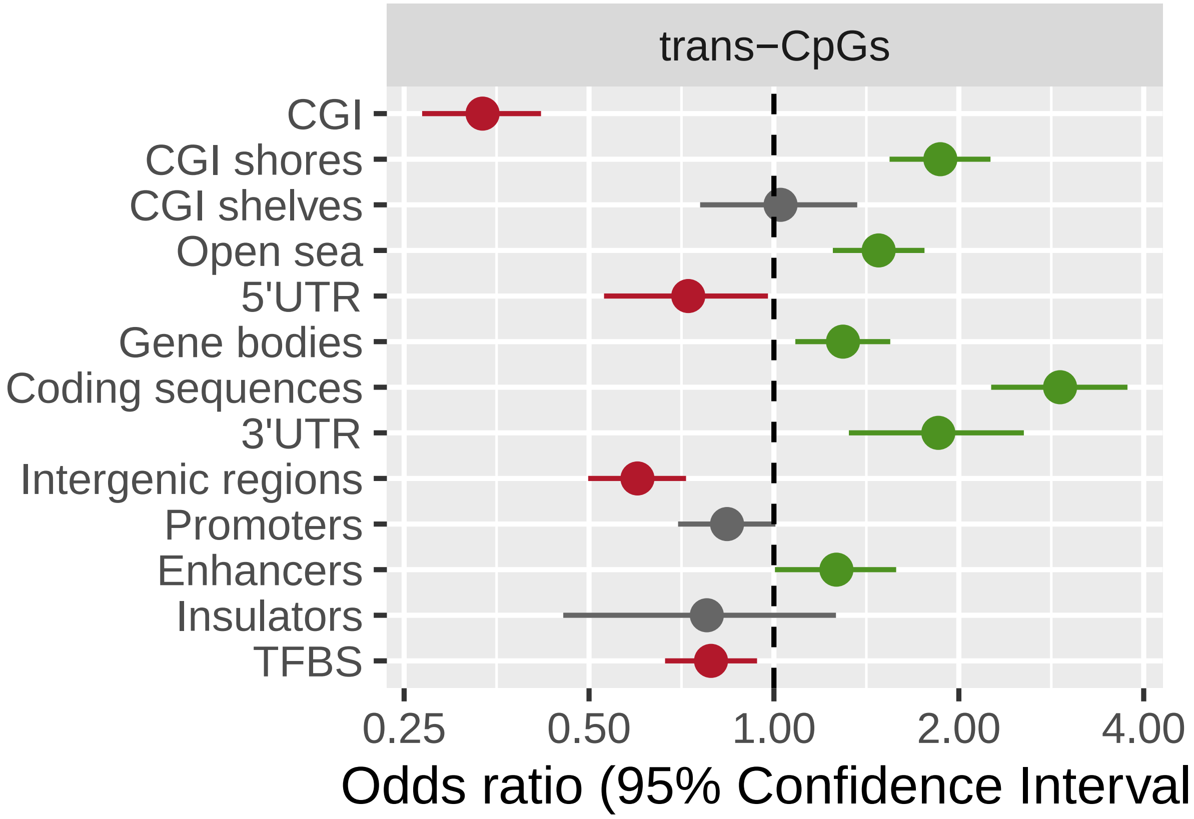

**Supplementary figure 17. Enrichment in genomic annotations of CpGs associated with *trans*-meQTL SNPs in the MHC.** The $x$-axis indicates the odds ratio and its 95% confidence interval (in logarithmic scale) for CpGs located within a specific genomic annotation. Background CpGs for comparison were the full set of CpGs with meQTLs. Significant enrichment is marked in green, depletion in red, and non-significant genomic annotations in grey.

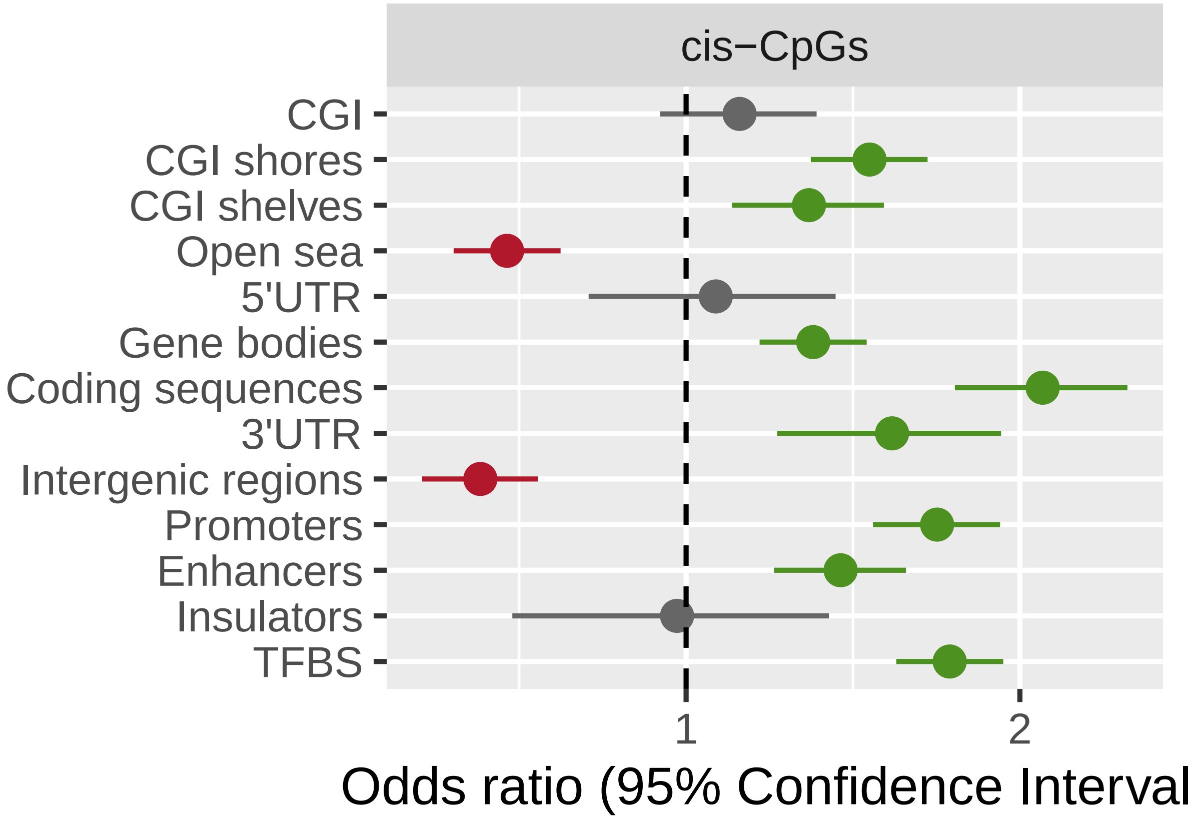

**Supplementary figure 18. Enrichment in genomic annotations of CpGs associated with complex phenotypes after SMR.** The $x$-axis indicates the odds ratio and its 95% confidence interval (in logarithmic scale) for CpGs with cis-meQTL co-localised with GWAS signals of complex phenotypes, located within a specific genomic annotation. Background CpGs for comparison were the full set of CpGs with meQTLs. Significant enrichment is marked in green, depletion in red, and non-significant genomic annotations in grey.
